# SufB intein splicing in *Mycobacterium tuberculosis* is influenced by two remote conserved N-extein Histidines

**DOI:** 10.1101/2021.07.07.451452

**Authors:** Sunita Panda, Ananya Nanda, Nilanjan Sahu, Deepak Ojha, Biswaranjan Pradhan, Anjali Rai, Amol R. Suryawanshi, Nilesh Banavali, Sasmita Nayak

## Abstract

Inteins are auto-processing domains that implement a multi-step biochemical reaction termed protein splicing, marked by cleavage and formation of peptide bonds. They excise from a precursor protein, generating a functional protein via covalent bonding of flanking exteins. We report the kinetic study of splicing and cleavage reaction in [Fe-S] cluster assembly protein SufB from *Mycobacterium tuberculosis*. Although it follows a canonical intein splicing pathway, distinct features are added by extein residues present in the active site. Sequence analysis identified two conserved histidines in the N-extein region; His-5 and His-38. Kinetic analyses of His-5Ala and His-38Ala SufB mutants exhibited significant reductions in splicing and cleavage rates relative to the SufB wild-type precursor protein. Structural analysis and molecular dynamics simulations suggested that *Mtu* SufB displays a unique mechanism where two remote histidines work concurrently to facilitate N-terminal cleavage reaction. His-38 is stabilized by the solvent-exposed His-5, and can impact N-S acyl shift by direct interaction with the catalytic Cys1. Development of inteins as biotechnological tools or as pathogen specific novel antimicrobial targets requires a more complete understanding of such unexpected roles of conserved extein residues in protein splicing.

## 1. Introduction

Protein splicing is a self-catalyzed event that generates a continuous extein protein by ligating two intein separated extein regions with a peptide bond. This post-translational auto-excision of the intervening intein protein is critical for the formation of an active protein(1–4). Splicing and cleavage products (Figure 1C) are obtained through a series of nucleophilic displacement reactions mediated in a coordinated fashion by the catalytic residues(5–9). Control over this reversible interruption of the functional form of the host exteins can play a regulatory role in protein activation (10, 11). The extein sequences upstream and downstream of N-terminal and C-terminal intein ends are termed ‘N-extein’ and ‘C-extein’ respectively. The intein folds into a horseshoe-shaped structure with a catalytic cleft that brings the conserved catalytic residues and the extein splice junction close enough for initiation of splicing reaction(12).

**Figure 1.**
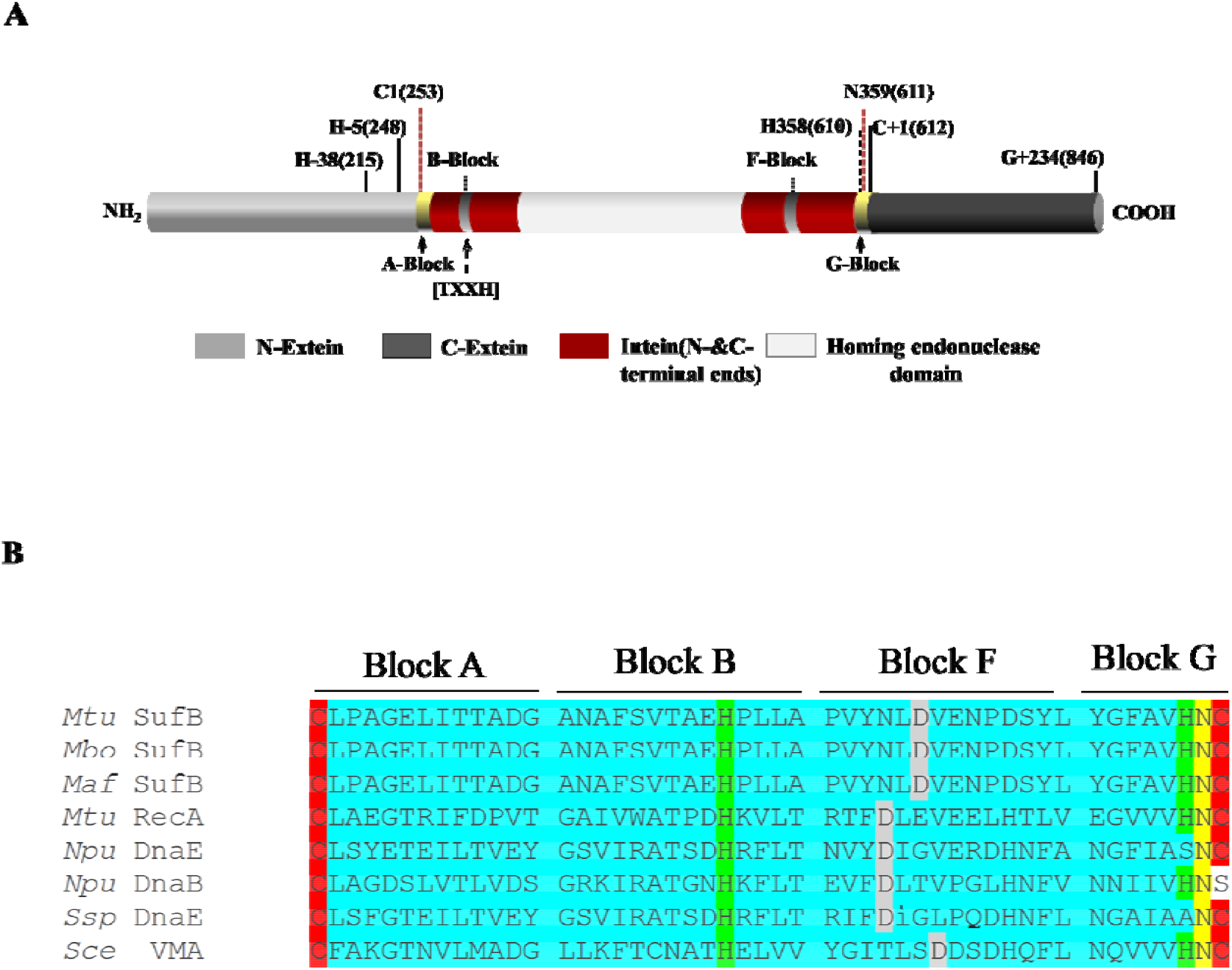
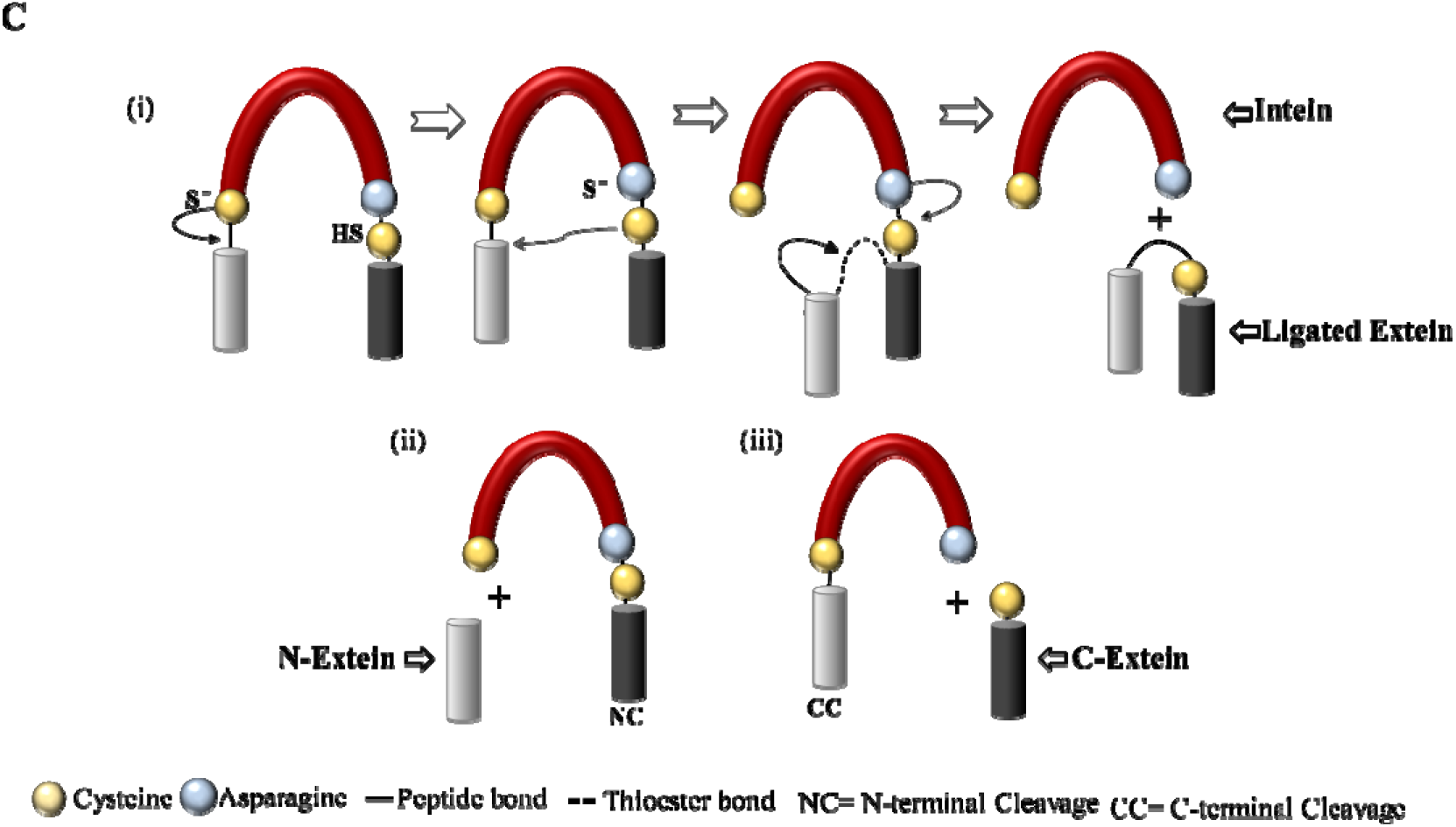
(A) Structural domains of the *Mtu* FL-SufB precursor. Important residues for the intein (numbered 1, 2, …), N-extein (numbered -1, -2, …) and C-extein (numbered +1, +2, …) are shown at the top. The conserved N-extein histidines are H-5 (H248 in *Mtu* FL-SufB) and H-38 (H215 in *Mtu* Fl-SufB). (B) Multiple sequence alignment of *Mtu* SufB with other intein-containing precursors shows conserved residues (highlighted in different colors) that participat in a canonical splicing pathway. (C) (i)Mechanism of intein splicing comprising of sequential nucleophilic attacks by C1 and C+1 to generate a branched intermediate by trans-esterification, cyclization of terminal Asn to resolve the branched intermediate ligates exteins and to cleave the SufB intein. (ii) N-terminal and (iii) C-terminal cleavage to form off-pathway products.

Inteins themselves have N-terminal and C-terminal regions with conserved sequence segments (termed as Blocks or motifs) that facilitates the splicing and cleavage reaction(s). N-terminal intein region comprises of A, N2, B, and N4 structural motifs or Blocks while F- and G-Blocks are part of the C-terminal intein region (13). Typically, A-Block contains Cys/Ser or Thr; B-Block includes His and Thr residues; F-Block usually has Asp and His and the G-Block bears two conserved residues; a penultimate His and a terminal Asn (8, 14, 15). Classical or Canonical (Class 1) intein splicing involves 4 sequential acyl rearrangements (Figure 1C), a nucleophilic attack by C1 or S1/T1 leads to an N-S/N-O acyl shift converting the peptide bond of N-terminal splice junction to a thioester linkage, a second nucleophilic attack by C+1 forms a branched intermediate at C-terminal splice junction through esterification, the branched intermediate is resolved by terminal Asn cyclization through cleavage of C-terminal splice junction, and finally an (S-N/O-N) acyl shift fully ligates the two extein segments by an amide bond formation (16–18).

These bond rearrangements at splice junctions during the cleavage and splicing reaction(s) are assisted by non-catalytic intein residues through stabilization of various intermediate structure(s)(19).The Block A residues Cys, Ser, or Thr participate in the first step of splicing with significant assistance from the Block B His and Thr residues. The highly conserved Block B His destabilizes the scissile peptide bond either by reducing the energy barrier or by loss of resonance or protonation of the Cys1 amide bond via His imidazole ring to catalyze the N-S acyl shift (20–24).The Block F Asp residue drives the thio-esterification through the tetrahedral intermediate by ground-state destabilization (8, 22, 25). It is also proposed that Block B histidine plays a dual catalytic role; being weakly basic it deprotonates the Cys1 to accelerate the N-S acyl shift and subsequently acts as an acid to stabilize the tetrahedral intermediate(21, 23, 24). The Block F Asp residue also deprotonates the C+1 residue to stabilizes the net positive charge on Cys1 that drives the transesterification reaction. The F- and G-Block His residues are critical in the coordination of terminal Asn cyclization. The F-Block His increases nucleophilicity of Asn, and the G-Block His accelerates the Asn cyclization by increasing the electrophilicity of backbone peptide (26–29). The final acyl-shift is energetically favorable and does not require assistance from either intein or extein residues(30). Inteins do show polymorphisms in the catalytic residues leading to variation in the splicing mechanism as seen in Class 2 and Class 3 intein splicing (11, 31–35).

Interactions between extein residues and the catalytic intein core in the regulation of splicing reaction have been studied by modulating intein activity, changing the conformation of catalytic cleft, and restraining the activity of catalytic residues (6, 15, 36–39). Previous studies on intein-extein partnership in intein splicing have suggested mediation by extein residues both near and remote to the N- and C-terminal splice junctions (7, 15, 36, 38, 40–42).The N-extein residue at the first position (−1) is important for the first thioester reaction and shows enhanced N-terminal cleavage rate (>4-fold) or attenuated cleavage by 1,000-fold by replacing the native residue to aspartate and proline respectively (36). Substitution of bulky amino acids at this position can cause local distortion and induce N-cleavage reaction (23). The participation of the first C-extein Cys+1 in the second and third steps of the splicing reaction has been demonstrated by its mutation dramatically augmenting or inhibiting splicing and generating off-pathway N-cleavage products(12, 14, 17, 40, 43). Earlier, extein effects were assumed to be limited to residues proximal to the intein (12, 15, 44), but recent work has shown that distal exteins are implicated as environmental sensors with the role in regulating splicing depending on solution environment and temperature in *Pho* RadA precursors (6, 10, 39).

SufB is a critical component of [Fe-S] cluster assembly and repair machinery called SUF (mobilization of sulfur) complex. This is a stress response system that gets upregulated during periods of oxidative stress and Fe starvation (45, 46). Though there are multiple pathways for [Fe-S] cluster biogenesis among the three kingdoms of life such as NIF (nitrogen fixation) and ISC (Iron-sulfur cluster), the SUF complex is unique in mycobacteria. Fe-S cluster containing proteins execute a broad spectrum of cellular functions in organisms such as respiration, gene regulation, RNA modification, DNA repair, and replication (47–49). Although the SUF system has been well characterized in the *E. coli* system,(17, 45, 46, 50–52), the importance of SufB intein splicing in the formation of functional Suf complex has been shown in mycobacteria (52).

The present study reports on the splicing and cleavage reactions of a full-length *Mtu* SufB (FL-SufB) precursor protein. We delineated the different structural domains of *Mtu* SufB, analyzed whether it follows a canonical or non-canonical intein splicing pathway, identified intein and extein residues that participate in catalytic cleft formation, assessed both their conservation in different mycobacterial species and their role in regulating cleavage and/or splicing reactions, and analyzed distinctions from other intein precursor proteins. We found intein residues highly conserved in different mycobacterial species that favor a canonical splicing mechanism (Figure 1A and 1B). We detected two distal histidines in the N-extein region, His-5 and His-38, that are conserved in mycobacteria, archaea, and other microbes where SUF is the exclusive system for Fe-S cluster biogenesis (Figure 2A and 2B). Biochemical analyses of H-5A and H-38A SufB mutants confirmed their influence on splicing and cleavage. Structural modeling of *Mtu* SufB and explicit-solvent molecular dynamics (MD) simulations of the model were used to analyze the SufB precursor splicing active site dynamics, and these simulations suggested that N-terminal cleavage could be supported by an interaction between H-38 and H-5. These observations suggest that the two distal H-5 and H-38 N-extein residues participate in SufB precursor stabilization and aid *Mtu* SufB intein splicing.

**Figure 2.**
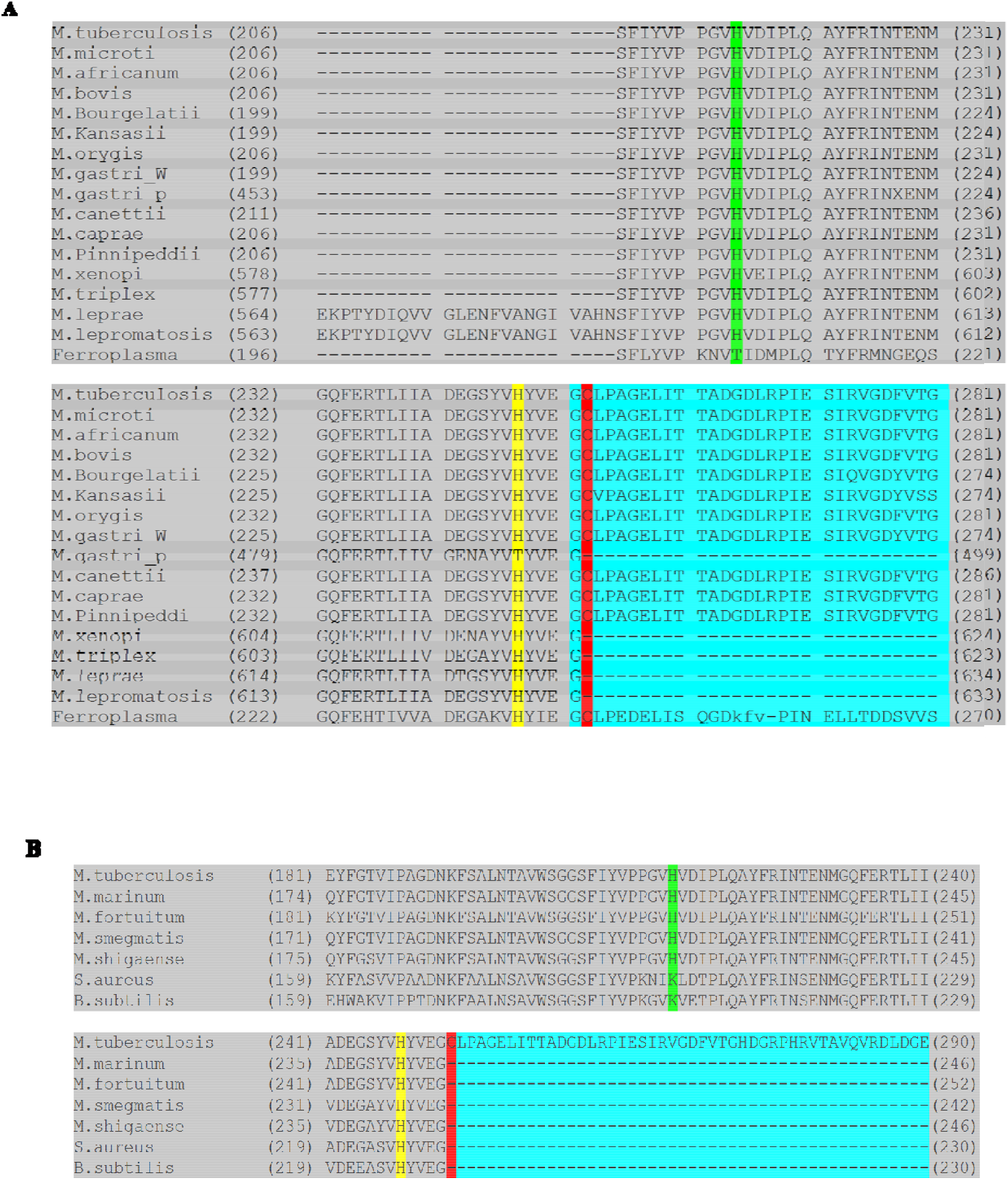
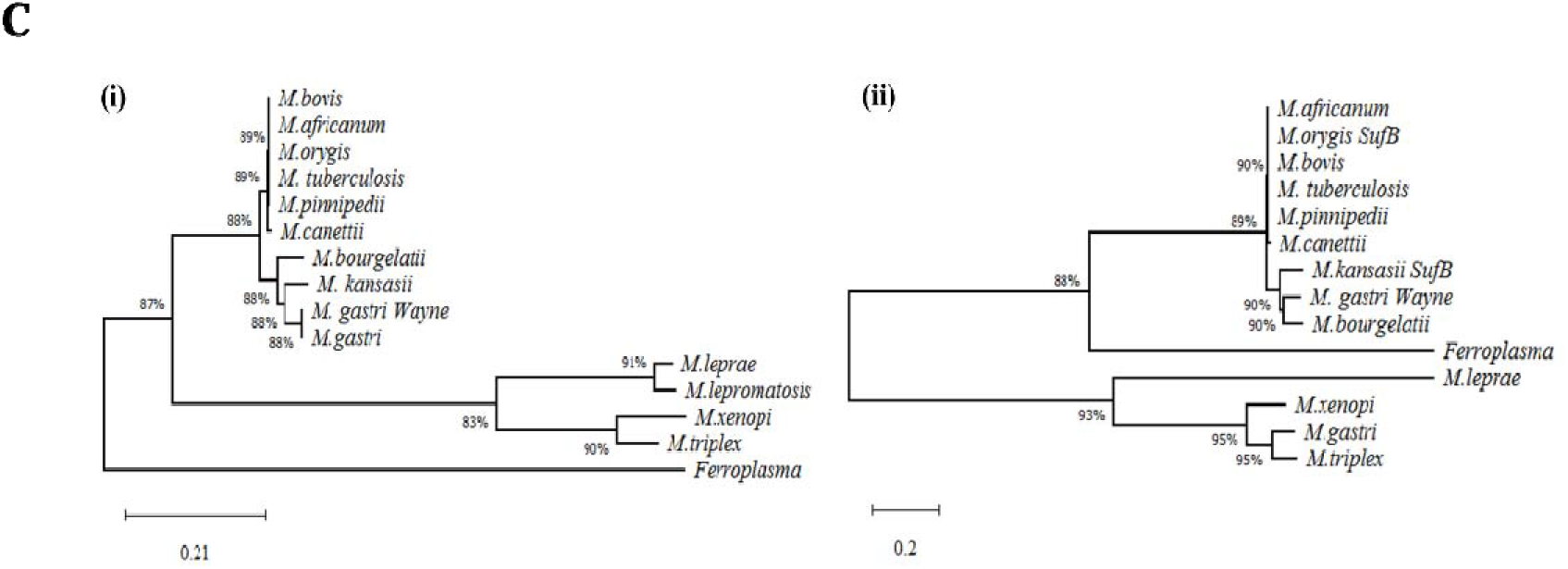
Conserved histidines in the *Mtu* FL-SufB N-extein. (A) Multiple sequenc alignment focused on the junction between the N-extein and intein in mycobacterial and archaeal species SufB proteins with conserved H-5 (yellow), H-38 (green), C1 in intein (red). N-extein and intein residues are delineated by shading in grey and cyan blue, (B) Multiple sequence alignment focusing on N-extein∼intein junction in mycobacteria and organisms where SUF is the exclusive system for Fe-S cluster assembly, (C) phylogenetic analyses of (i) full-length (FL) SufB protein and (ii) SufB intein sequences in mycobacteria and *Ferroplasma acidarmans*.

Kinetic analyses of H-5A SufB mutant demonstrated 3-fold and 1.4-fold reductions in splicing and cleavage rates respectively, relative to wild-type *Mtu* SufB precursor. Likewise, a 3.4-fold and 3-fold reduction in splicing and cleavage rates were observed in the H-38A SufB mutant. A side-by-side *Mtu* SufB structure prediction was done using homology (chimera) modeling, secondary structure prediction through consensus with other protein sequences. Subsequently, molecular dynamics (MD) simulations in aqueous media were carried out to find the equilibrated structure. Furthermore, MD simulations clarified the structural features of the SufB intein active site and indicated that N-terminal cleavage reaction is catalyzed by H-38 with the assistance of H-5. Taken together, our study substantiates a distinct mechanism for N-terminal cleavage reaction shown by *Mtu* SufB. Although H-38 is relatively distal to the N-terminal splice junction when supported by His-5, it can efficiently activate the first step of splicing reaction. Finally, we have proposed a novel mechanism for the N-cleavage reaction mediated via the concerted actions of these conserved histidines in the N-extein region of *Mtu* FL-SufB precursor. These observations suggest that H-5 and H-38 might have important biological role(s) in the SufB precursor stabilization and perhaps the functionality of *Mtu* SufB protein.

## 2. MATERIALS AND METHODS

### 2.1. Genetic constructs

The full-length Fe-S cluster assembly protein SufB from the *Mycobacterium tuberculosis* H37Rv strain (*Mtu* FL-SufB) and its isolated intein (*Mtu*-SufB-I) genes were PCR-amplified using Pfu Ultra High-Fidelity DNA Polymerase (Agilent Technologies) from heat-killed *Mtu* genomic DNA. DNA purification by gel electrophoresis was followed by EcoRI and HindIII restriction digestion and cohesive end ligation (T4 DNA Ligase, NEB Cat. No. M0202S) for cloning. The genes were inserted into the multiple cloning site 1 (MCS1) of the low copy expression vector pACYCDuetTM-1(Novagen), which was driven by a T7 promoter/lac operator with a chloramphenicol resistance gene for selection. The constructs were screened via colony PCR and confirmed by sequencing (Sequencing Core Facility, SUNY, Albany, and Eaton Bioscience Inc. sequencing service) using the ACYCDuetUP1 (Novagen Cat. No.71178-3) and DuetDOWN1 primers (Novagen Cat. No. 71179-3), as well as the original primers. *Mtu* FL-SufB mutants H5A, H38A, C1A, N359A were generated by substituting respective key catalytic residues to alanine via phosphorylated inverse PCR primers. Splicing inactive (SI) *Mtu* FL-SufB double mutant (C1A/N359A) was created via inverse PCR to add the N359A mutation into the C1A cleavage mutant. The cloned H-38A SufB mutant genes were also confirmed separately by sequencing (Agrogenomics, Odisha) using ACYCDuetUP1 (IDT Cat. No. 103948189) and DuetDOWN1 (IDT Cat. No. 103948190) along with the SufB Primers. All the above primers are listed in Table S1 (Supplementary materials).

### 2.2. Sequence analysis

Protein sequences for *Mtu* FL-SufB (Accession number YP_006514844.1, GI: 397673309) and the 477 amino acid intein-less SufB protein from *Mycobacterium smegmatis* strain MC2 155 (*Msm*-SufB-FL, accession number YP_887437.1, GI: 118472504) were pairwise sequence aligned in ClustalW to distinguish extein and intein regions in *Mtu* FL*-*SufB (53). The identified *Mtu* SufB intein sequence (*Mtu* SufB-I) was confirmed by sequence comparison using Blast with the sequences deposited in Inbase (The Intein Database, www.inteins.com) (54). Different structural domains of the intein-like homing endonuclease, and the N- and C-terminal inteins were clearly demarcated by sequence alignment and structural analysis of SufB intein with homing endonuclease domain (I-CreI) and intein homing endonuclease Ii (PDB: 2CW7). The *Mtu* SufB-FL and *Msm*-SufB-FL sequences, combined with one archaeal and other bacterial SufB proteins collected from the NCBI and intein databases (54), were aligned using Dialign2 software (55). Conservation of different intein and extein residues was edited and color-coded manually. Phylogenetic tree analysis was performed using the Maximum likelihood method in the MEGA X program (56, 57) for both SufB inteins [Figure 2C (ii)] and SufB precursor sequences [Figure 2C (i)] from different organisms.

### 2.3. Protein overexpression and purification

Full-length (FL) un-spliced precursor and mutant SufB proteins carrying an N-terminal 6X-His-tag were over-expressed in BL21 (DE3) *E.coli* cells via IPTG (500µM) (Sigma 367-93-1) induction at 37°C for 4hrs. Cells were resuspended in lysis buffer (20mM sodium phosphate, 0.5M NaCl, pH 7.4) and lysed via tip sonicator (Sonics vibra cell VCX-130). Proteins were over-expressed and isolated from inclusion bodies (IB) via centrifugation. The IB materials were solubilized by 8M urea (23) (Merck, 1084870500) buffer (lysis buffer, 8M urea, 20 mM of imidazole (MP–biochemicals-288-32-4) and centrifuged at 16,500g for 20 min to collect the supernatant. Then 6X-His-tagged wild type SufB precursor and mutant proteins were purified by Ni-NTA affinity column (Ni-NTA His trap, HP GE Healthcare Life Sciences-17524802) (23, 58–62). Prior to sample application, columns were equilibrated with binding buffer (20mM sodium phosphate, 0.5M NaCl, 40mM imidazole). After sample loading, columns were washed several times (15 CV) in binding buffer. Finally, proteins were eluted as purified fractions in elution buffer (20mM sodium phosphate, 0.5M NaCl, 500mM imidazole) followed by quantification via Bradford’s assay.

### 2.4. *In-vitro* splicing and cleavage assays

2.5 µM of purified proteins were allowed to refold in 1ml of renaturation buffer (20 mM sodium phosphate, 0.5 M NaCl, 0.5 M Arginine,1 mM EDTA, pH 7.4) in presence of 2mM TCEP-HCl (sigma-51805-45-9) at 20°C for 24h (58–63). The 0hr sample was retrieved before renaturation and splicing was quenched by addition of loading dye (0.1% bromophenol blue, 50% glycerol, ß-mercaptoethanol, 10% SDS, tris 6.8) followed by rapid freezing at -20°C. Our controls, splicing inactive(SI) SufB double mutant and empty expression vector pACYC Duet-1, were treated similarly for the *in-vitro* assays. For the N-cleavage assay, proteins were refolded in presence of reducing agents and nucleophiles such as 2mM TCEP-HCl, 50 mM DTT (Roche-10708984001), and 0.5 M 250 Hydroxylamine (SRL-66164) with 1mM TCEP in renaturation buffer (58–63). For splicing and cleavage analysis, sample extraction at each time interval was followed by the addition of loading dye to stop the reactions and then boiling at 95°C for 5 min. Resultant products from various refolding reactions were resolved through 4∼10% gradient SDS PAGE. Protein bands were stained with Coomassie blue R-250 and densitometric analysis was performed by using GelQuant.Net biochemical solutions. Percentage(s) of splicing and cleavage products were measured by taking the percentage(s) of the ratio of the total splicing product (LE and I) over total proteins (LE+I+P) and total N-cleavage product (NE+NC) over total proteins (NE+NC+P). The 0hr splicing value(s) were subtracted at each time point for baseline correction.

### 2.5. Kinetic analyses

Since *Mtu* 6X (His)-tagged WT SufB and SufB mutants (H-5A, H-38A, and SI C1A/N359A) were purified and renatured at different temperatures, after normalizing splicing and cleavage values at different time intervals, the plot was generated by taking the percentage of splicing or cleavage product with respect to time (in min). Next, the curve was fitted in pseudo-first-order kinetics, with an equation Y=Y0 + (Plateau-Y0) *(1-exp(-K*x)) [Where X=time, Y0 =Y value when time (X)= time0, Plateau= max Y value at time t, K = rate constant, expressed in reciprocal of the X-axis (time units)] in graph pad prism software. The fitted curve was generated by automatic outlier elimination fitting in a nonlinear regression equation. The rate constant (K) and Vmax were generated by the software. Half-life (t1/2) was calculated by graph pad prism using the formula (Ln2/K).

### 2.6. Western blot

Western blot analysis was performed using an anti-His antibody (Invitrogen, LOT 1902132) to confirm the identity of splicing and cleavage products. Following resolution through SDS PAGE, test proteins were transferred to a nitrocellulose membrane; at 50v, 2h. After a successful transfer, blocking was done with 5% skim milk for 2h at room temperature. Then the blot was incubated with HRP conjugated anti-His antibody (Invitrogen, LOT 1902132) at 1:5000 dilutions for 16h at 40 C. Then blot was washed with 1X TBST and developed using 13 ECL as the substrate. N-extein detection was done with 1:2500 antibody dilution.

### 2.7 Mass spectrometry and chromatography

After renaturation, proteins were resolved through 4∼10% SDS PAGE. Protein identification by mass spectrometry was performed at Central Proteomics Facility, Institute of Life Sciences, Bhubaneswar. In gel digestion of protein and extraction followed the protocol described by Shevchenko et. al. (64). Extracted peptides were desalted by using C18 Zip-Tip (Millipore) and then dried in a centrifugal vacuum concentrator. Peptide samples were reconstituted with 0.1% TFA in 50% ACN solution, mixed (1:1) with α-cyano-4-hydroxycinnamic acid solution (C8982; 10 mg/ml) and then spotted onto metal target plate in triplicates. MS and MS/MS spectra were acquired using MALDI TOF/TOF (AbSciex TOF/TOF 5800). Acquired spectra were searched against *Mycobacterium tuberculosis* from NCBInr database using Mascot search algorithm (version 2.0; Matrix Science, Boston, Massachusetts) and protein pilot software (SCIEX, USA) for protein identification (65, 66). Searches were performed allowing trypsin mis-cleavage up to 1; Carbamido-methylation of cysteine and oxidation of methionine as variable and fixed modification respectively. The peptide mass tolerance was set as 100 ppm for precursor ion and 0.8 Da for fragment ion with +1 charge. Since all the splicing and cleavage products were identified as *Mtu* FL-SufB, the acquired spectra were researched against a customized database containing probable splicing products and cleavage products of SufB protein of *Mycobacterium tuberculosis* using mascot search engine in Protein Pilot Software with the same parameter. β and BSA were run as internal calibration. The protein score, percent coverage, theoretical molecular weight, and Iso-electric pH value were obtained. The mass spectrometry proteomics data have been deposited to the ProteomeXchange Consortium via the PRIDE [1] partner repository with the dataset identifier PXD015199.

Purified and renatured *Mtu* FL-SufB protein was examined by analytical HPLC carried out on an Agilent 1200 series instrument equipped with a Zorbax gf-450 column (6µm, 9.4×250mm) with a flow rate of 1ml/min. All HPLC runs used the following solvent: 1:1 water and isopropanol (solvent 1) and 0.1% TFA (Trifluoroacetic acid)in water (solvent 2). The column was equilibrated using solvent 1 and solvent 2. A protein sample was inserted into the column and ran for 30mins. The retention time vs protein intensity measured at 280 nm was noted. For reference, we ran 6µl of Precision plus protein ladder (Biorad1610374) diluted to 30µl using Sodium phosphate buffer. The retention time vs molecular weight of the known protein standards was measured and plotted to make a standard curve. The unknown protein peaks from the test samples were compared with the standard curve to find out the relative molecular weights. The molecular weights of the expected fragments from the MALDI-TOF/TOF MS (Table S3) data were compared with the standard curve and molecular weights of the unknown peaks were determined.

### 2.8 Molecular Dynamics (MD) Simulations

#### 2.8.1. Homology modeling of Mtu SufB precursor

A homology model of the *Mtu* FL-SufB precursor was built using the SufB chain of the *E. coli* SufB-SufC-SufD complex (PDB ID: 5AWF, chain A), the *S. cerevisiae* intein homing endonuclease (PDB ID: 1VDE, chain A), and the *T. kodakarensis* homing endonuclease (PDB ID: 2CW7, chain A) using Bioluminate 2.7 from Schro dinger. The model was assessed by Ramachandran plot analysis and PROSA(Z-score) (67).

#### 2.8.2. MD simulation analysis

Explicit-solvent MD simulations were performed using GROMACS (V5.1.4) (68) with the OPLS-AA force field (69) and the SPC/E water model (70, 71). The *Mtu* SufB model (846 aa) was solvated in a cubic box with at least 1 nm distance between the protein and the edge of the box, and neutralized with NaCl. The system was minimized with target Fmax of no greater than 1000 kJ mol-1 349 nm-1 with steepest descent minimization with a spherical cut-off at 1 nm was imposed on all intermolecular interactions with verlet cut-off scheme(72, 73). The leap-frog algorithm with a timestep of 2fs (74) and a canonical NVT ensemble was used to run the simulation for 100 ns, with temperature maintained at 300K through velocity-rescale coupling and no temperature coupling. H-bonds were constrained using lincs with the order of 4 (75). The particle-mesh Ewald (PME) algorithm was used for implementing long-range electrostatic interactions with the grid dimension of 0.16nm and interpolation order of 4 (76). Histidine at positions 5 and 36 were mutated to alanine by homology modelling, and MD simulations were performed on these mutant proteins in an identical fashion. The MD simulation trajectories were analyzed with GROMACS (68) and PyMOL Molecular Graphics System, Version 1.2r3pre, Schrödinger, LLC, and plotted using Origin 8.0.

## 3. RESULTS

### 3.1 Structural domains of *Mtu* full-length (FL) SufB precursor

Pair-wise sequence alignment of the *Mtu*-FL-SufB and the intein-less *Msm*-FL-SufB protein sequences using Clustal-W could delineate the intein and extein boundaries for *Mtu*-FL-SufB due to the ∼95% sequence similarity between the extein sequences. The boundaries of the 359 amino acid *Mtu* SufB (*Mtu* Pps1) intein from *M. tuberculosis* strain H37Rv were also affirmed by a Blast search of Inbase. The final demarcation of these structural domains is shown in Table S2 and is illustrated in Figure 1A. Multiple sequence alignment with other inteins shows conservation of catalytic cysteines (C1 and C+1), Block B His67, Block F Asp77, penultimate His358, and terminal Asn359 (Figure 1B). The presence of these conserved residues suggests that the *Mtu*-FL-SufB precursor (846 aa) is auto-processed by a canonical intein splicing pathway to ligate its N-extein (252 aa) and C-extein (235 aa) to form the native SufB protein (487 aa). The major by-product is the SufB intein (359 aa) containing an intein domain (155 aa) and an endonuclease domain (204 aa), with minor off-pathway products possible due to N-terminal cleavage (594 aa) and C-terminal cleavage (611 aa) (54, 77).

### 3.2 Conservation of His-5 and His-38 in different bacterial species

SufB from different mycobacterial and archaeal species exhibit high sequence similarity (Figure 2A and B). Phylogenetic analysis suggests that these mycobacterial proteins have a common ancestral origin (78). Cladogram analysis of SufB intein sequences from different mycobacterial species shows significant similarities, except in *M. leprae*, *M. lepromatosis*, *M. triplex* and *M. xenopi*, possibly due to different intein insertion sites (17). In Ferroplasma, which belongs to a different kingdom, there is divergence for both the intein [Figure 2C (ii)] and the full precursor [Figure 2C (i)]. These variations in the extein sequences and intein insertion sites suggests some divergent intein evolution and independent intein transfer in different species and kingdoms.

Two conserved histidines were identified in the N-extein region of different mycobacterial and archaeal SufB precursor proteins: His-5 and His-38 (Figure 2A). His-5 is conserved in intein-les SufB proteins from *Staphylococcus aureus*, *Bacillus subtilis*, and certain mycobacteria that use SUF complex as the sole pathway for [Fe-S] cluster generation (Figure 2B) (79, 80). His-38 is conserved in both intein-bearing and intein-less mycobacterial SufB proteins where the SUF system is required to synthesize [Fe-S] clusters (Figure 2B). Identification of two highly conserved metal-chelating residues proximal to N-terminal extein∼intein junction raises the possibility of their regulatory roles on cleavage and/or splicing as well as in the functionality of [Fe-S] cluster assembly protein SufB. These two histidines could participate in protein splicing either via direct or indirect interaction with the catalytic residues near cleavage site(s).

### 3.3 Kinetic study to evaluate the roles of conserved His-5 and His-38 on *Mtu* SufB intein splicing

H-5A and H-38A single mutants were generated to test for a regulatory effect of His-5 and His-38 on *Mtu* SufB intein cleavage and/or splicing. Such alanine substitutions for active site residues have previously resulted in a complete blockage of cleavage and splicing reactions (36). The splicing inactive (SI) SufB double mutant, with both C1 and N359 mutated to alanine (C1A/N359A), which is expected to abolish intein splicing completely (81), was used as a negative control.

Remnant 6x(His)-tagged *Mtu* FL-SufB precursor (P, 95.98 KDa), N-terminal cleavage (NC, 65.26 KDa) product, N-extein (NE, 29.9 KDa), ligated exteins (LE, 55.7 KDa), and intein (I, 40.2 KDa) were all observed upon *in vitro* refolding followed by SDS PAGE, and western blot analysis of the splicing of the *Mtu* FL-SufB precursor and its H-5A and H-38A mutants (Figure 3). As expected, none of the post-reaction products were seen in the case of SI double mutant (Figure S2A) and transformants expressing empty vector pACYC Duet-1 (Figure S2B). Possibly due to protein degradation, the C-terminal cleavage product (CC, 70.12 KDa) and C-extein (CE, 25.76 KDa) were not detected, which precluded analysis of the effects of H-5A and H-38A mutations on C-terminal cleavage.

**Figure 3.**
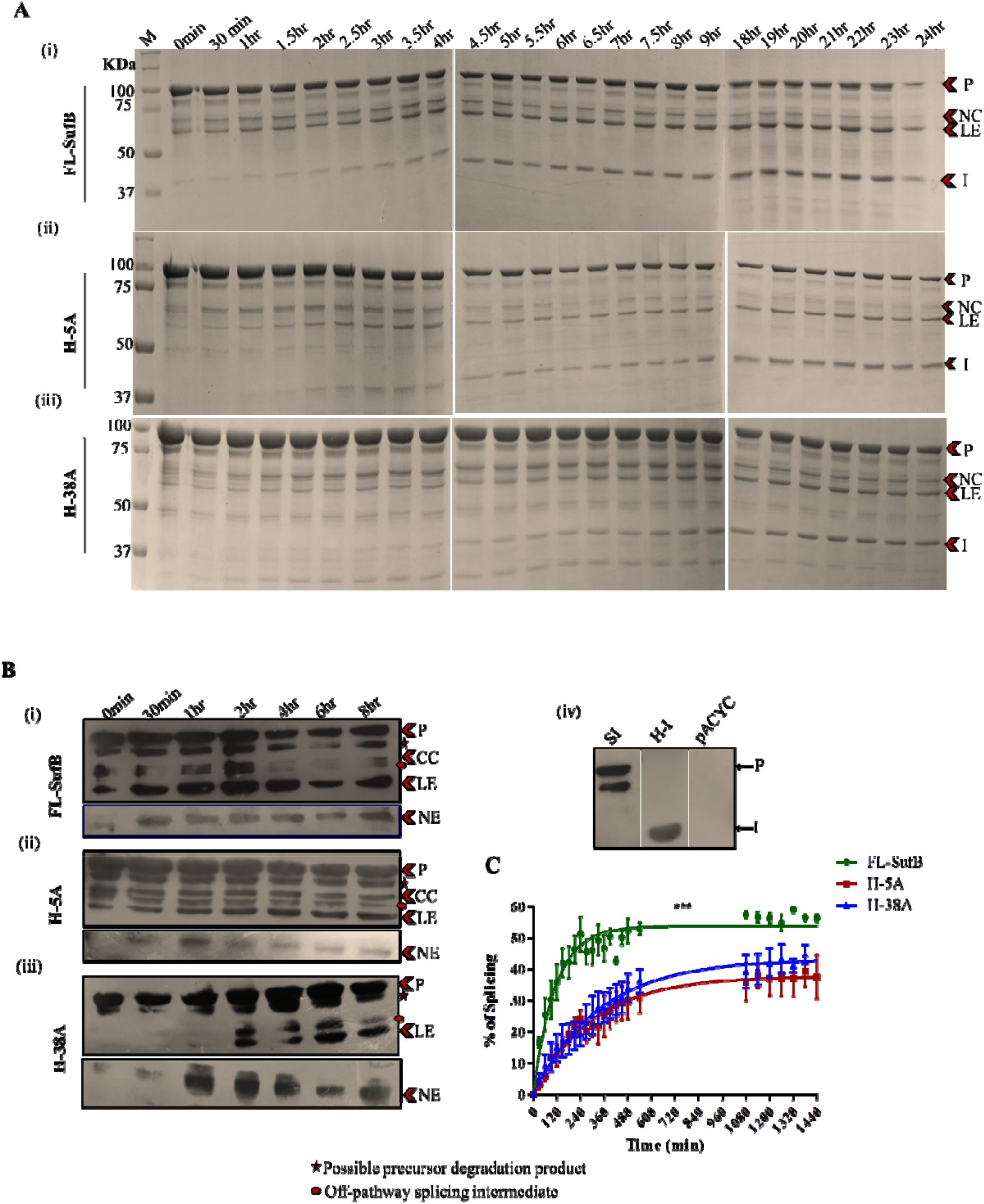
Effect of H-5A and H-38A mutations on *Mtu* SufB splicing. (A) Products from *in vitro* refolding reactions were resolved through 4∼10% gradient SDS PAGE for (i) *Mtu* FL-SufB precursor, (ii) H-5A, and (iii) H-38A mutant proteins exhibiting splicing over different time periods. (B) Western blot using anti-His antibodies confirm the identity of splicing and cleavage products for 6x(His)-tagged (i) *Mtu* FL-SufB precursor,(ii) H-5A,(iii) H-38A and (iv) the controls SI (splicing inactive double mutant SufB), H-I[6x(His)-tagged SufB intein], and pACYC Duet-1(cell lysates expressing empty expression vector). NE was blotted separately with higher concentration of primary antibody. (C) Splicing kinetics in *Mtu* FL-SufB precursor, H-5A, and H-38A with product quantities over different time periods fit to a pseudo first order reaction; Y=Y0 + (Plateau-Y0) *(1-exp(-K*x)).A statistically significant difference in splicing efficiency (p<0.0001) is observed between *Mtu* FL-SufB precursor, H-5A and H-38A mutants. Error bars represents (±1) SEM from 3 independent sets of experiments. P=Precursor, CC=C-cleavage, NC=N-cleavage, LE=Ligated Extein, I=Intein, NE=N-Extein, M= Protein marker.

#### 3.3.1 H-5A and H-38A mutants exhibit attenuated splicing reaction

Densitometric analysis was performed after *in vitro* renaturation of FL and mutant SufB proteins over a period for 24h. Precursor (P) proteins detected at time 0h were comparable for FL and mutant proteins (Figure 3A). Splicing efficiency was calculated as a percentage of splicing [(I+LE/P+I+LE) x 100] and plotted over time as pseudo-first-order reaction kinetics (Figure 3C). At 20°C, at least 3-fold reductions in splicing efficiency were observed for H-5A and H-38A, relative to *Mtu*-FL-SufB precursor (p<0.0001; one-way ANOVA, Table 1).

**Table 1.** Comparative analysis of different kinetic parameters for splicing in FL-SufB precursor, H-5A and H-38A SufB mutant proteins, at 20 °C temperature (data also shown in Figure 3C).

| Kinetic parameters | FL-SufB | H-5A | H-38A |
| --- | --- | --- | --- |
| $Y_{\max}$ | 53.81±1.03 | 37.82±1.68 | 43.35±1.7 |
| Rate constant (K)<br>(min<sup>-1</sup>) | 9.92±0.007$\times 10^{-3}$ | 3.3±0.0003$\times 10^{-3}$ | 2.9±0.0003$\times 10^{-3}$ |
| Half-life($t_{1/2}$) | 1.1h | 3.4h | 3.8h |

### 3.4 Kinetic studies of *Mtu* SufB N-terminal cleavage

The equilibrium of the first N/S acyl transfer step was examined using the reagents DTT (Dithiothreitol), and HA (Hydroxylamine), which induce N-terminal cleavage reaction. DTT can act both as a nucleophile and reducing agent. HA acts as a nucleophile and like DTT targets the linear thioester intermediate in the first step of splicing (82). The reducing agent Tris (2-carboxyethyl) phosphine (TCEP) was used as a control (83). N-cleavage reaction products were analyzed by SDS PAGE (Figure 4, 5, and Figure S3) and densitometric analysis of the cleavage products.

**Figure 4.**
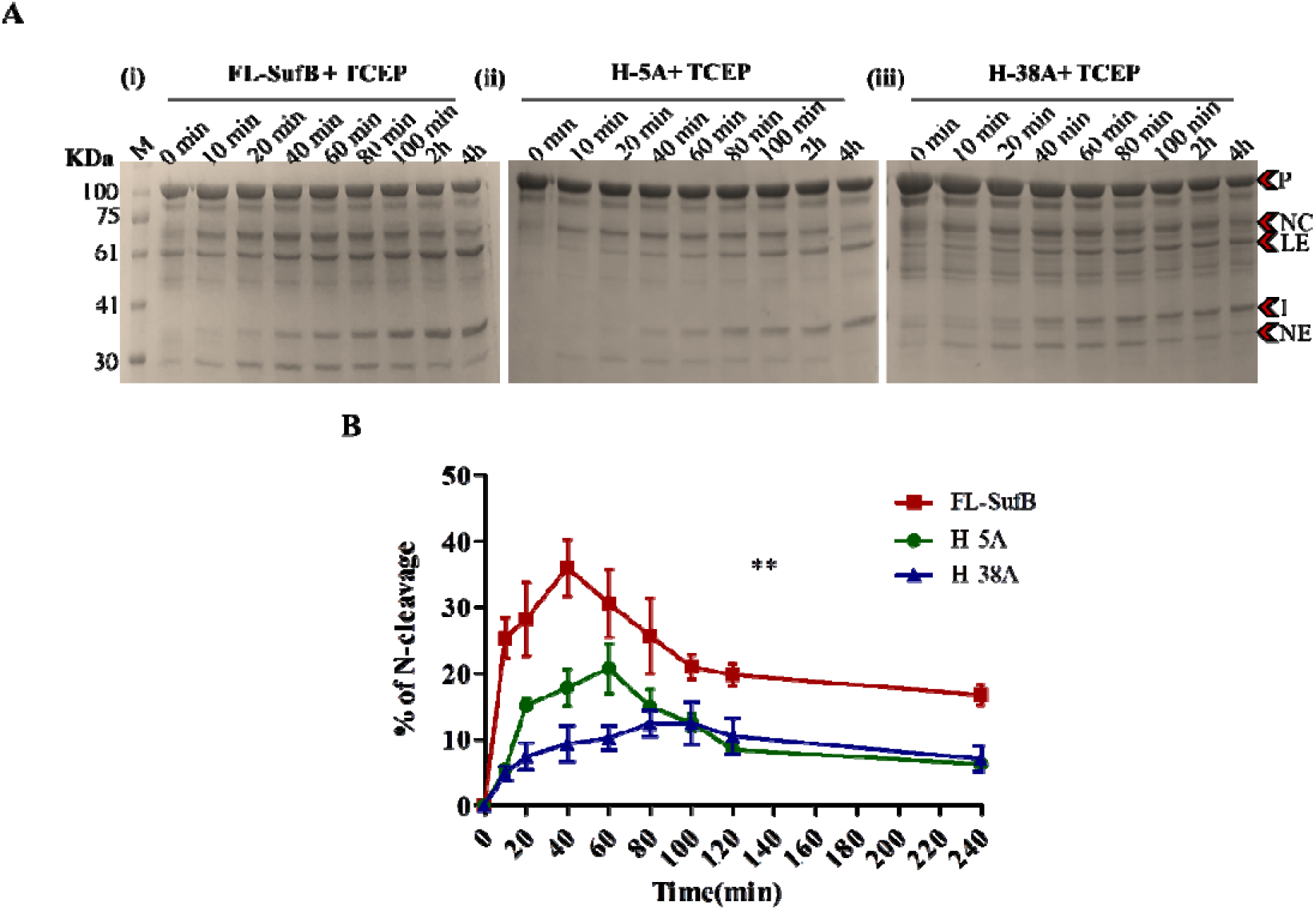
TCEP mediated N-cleavage in *Mtu* SufB: (A) SDS PAGE analysis of (i) *Mtu* FL-SufB precursor, (ii) H-5A and (iii) H-38A proteins displaying N-cleavage reactions over different time periods; (B) Statistically significant differences (P=0.0011, one-way ANOVA) are seen between TCEP mediated N-cleavage in *Mtu-*FL-SufB (i), H-5A (ii), and H-38A (iii). All the experiments were performed in triplicate and error bars represent (±1) SEM. P=Precursor, NC=N-cleavage, LE=Ligated Extein, I=Intein, NE=N-Extein, CC= C-Cleavage.

**Figure 5.**
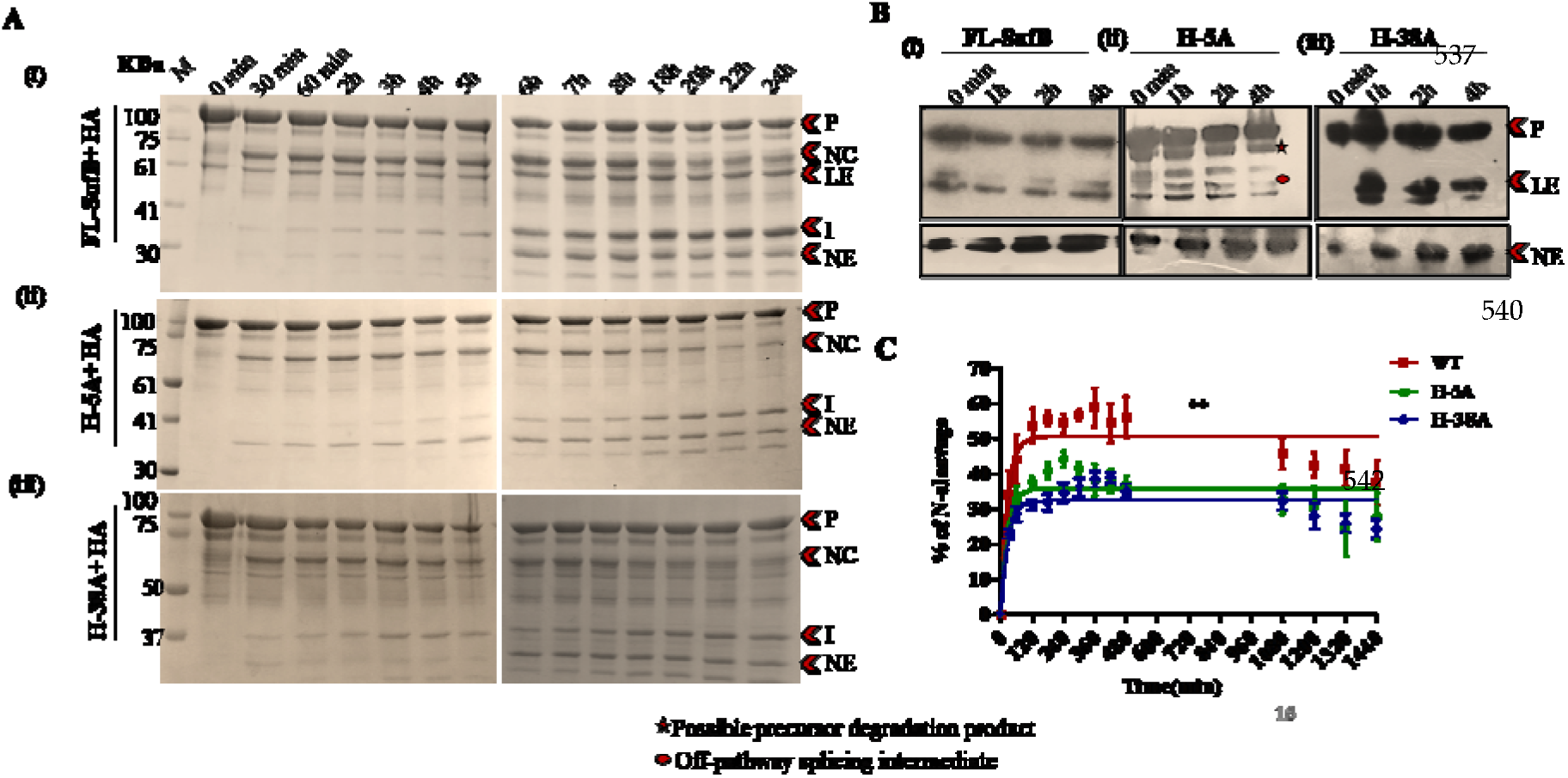
HA mediated N-cleavage in *Mtu* SufB: (A) SDS PAGE analysis of HA induced N-cleavage reactions in (i) *Mtu* FL-SufB precursor, (ii) H-5A, and (iii) H-38A; (B) Western blot analysis of splicing and cleavage products for HA induced N-cleavage reaction in (i) *Mtu* FL-SufB precursor, (ii) H-5A, and (iii) H-38A. NE is blotted separately with higher concentration of primary antibody; (C) Kinetic analysis of HA induced N-cleavage in WT, H-5A and H-38A mutant proteins suggests statistically significant differences (P= 0.0025; one way ANOVA) in % of N-cleavage. All the experiments were performed in triplicates and error bars represent (±1) SEM. P=Precursor, NC=N-cleavage, LE=Ligated Extein, I=Intein, NE=N-Extein, CC= C-Cleavage product.

#### 3.4.1 H-5A and H-38A mutations reduce TCEP mediated N-cleavage

The H-5A and H-38A mutant proteins showed diminished production of N-cleavage (NC) products relative to *Mtu* FL-SufB [Figure 4, Table 2(i)], although a proper fit for the reaction time course was not obtained through linear or non-linear regression. In *Mtu*-FL-SufB, N-cleavage was accelerated until 40 min followed by a decline as splicing became distinct. H-5A and H-38A mutants presented a sluggish course with the N-cleavage product peak at 60-80 min and then tapered off. At 40 min and 60 min, H-5A displayed 2-fold and 1.5-fold decrease in % N-cleavage respectively, relative to FL-SufB. Likewise, at 40 min and 60 min, H-38A mutant displayed a 3.8-fold and 3-fold reduction in % of N-cleavage respectively relative to FL-SufB (Figure 4).

**Table 2.**
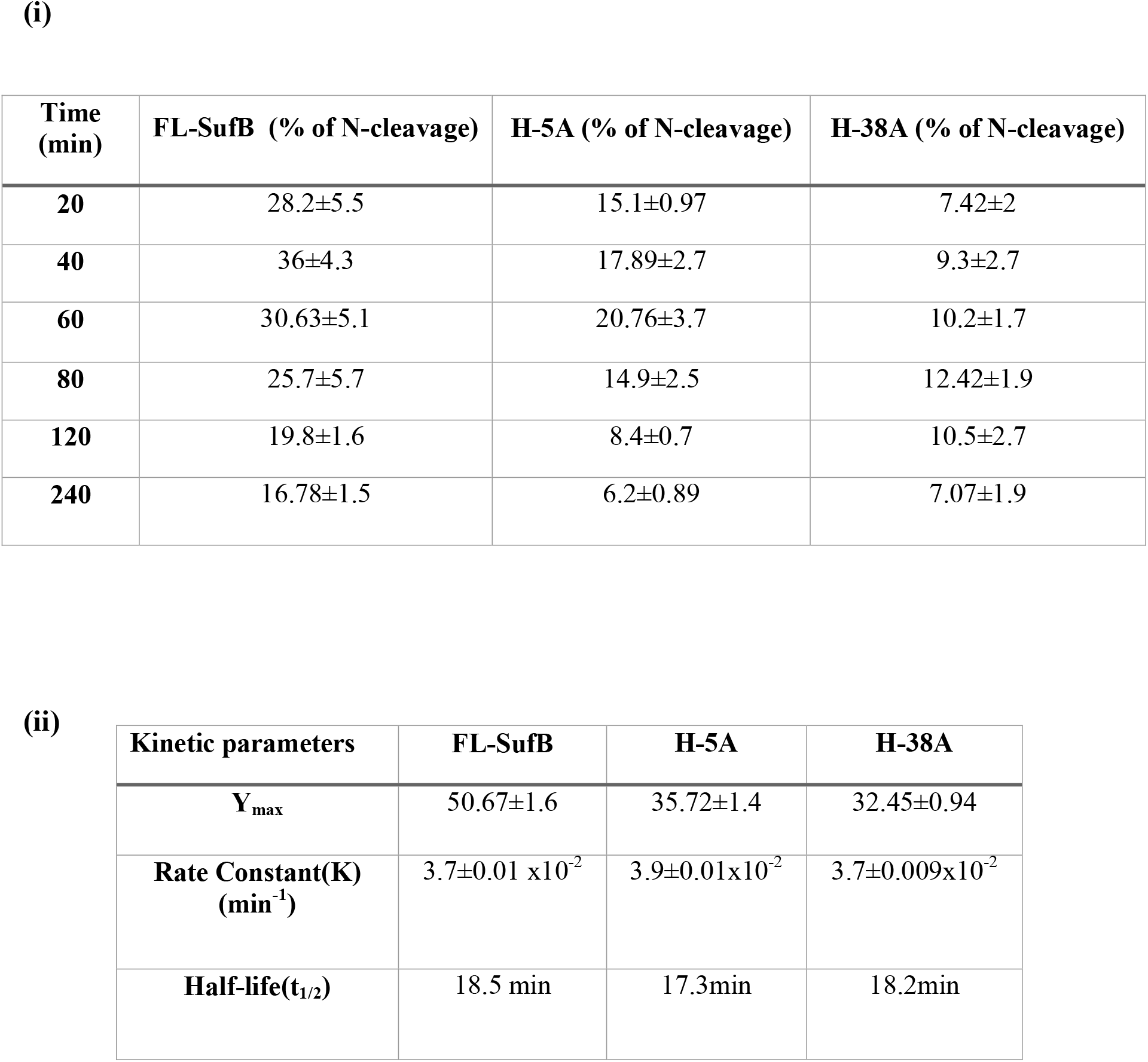
(i) Comparative analysis of TCEP mediated N-terminal cleavage reaction at 20 °C over 240 mins (data also shown in Figure 4B). (ii) Different kinetic parameters (Ymax, rate constant, and half-life) for HA-induced N-terminal cleavage in FL-SufB precursor, H-5A, and H-38A SufB mutants (data also shown in Figure 5C).

| Time (min) | FL-SufB (% of N-cleavage) | H-5A (% of N-cleavage) | H-38A (% of N-cleavage) |
| --- | --- | --- | --- |
| 20 | 28.2±5.5 | 15.1±0.97 | 7.42±2 |
| 40 | 36±4.3 | 17.89±2.7 | 9.3±2.7 |
| 60 | 30.63±5.1 | 20.76±3.7 | 10.2±1.7 |
| 80 | 25.7±5.7 | 14.9±2.5 | 12.42±1.9 |
| 120 | 19.8±1.6 | 8.4±0.7 | 10.5±2.7 |
| 240 | 16.78±1.5 | 6.2±0.89 | 7.07±1.9 |

(ii)
| Kinetic parameters | FL-SufB | H-5A | H-38A |
| --- | --- | --- | --- |
| $Y_{\max}$ | 50.67±1.6 | 35.72±1.4 | 32.45±0.94 |
| Rate Constant(K) (min<sup>-1</sup>) | 3.7±0.01 $\times 10^{-2}$ | 3.9±0.01$\times 10^{-2}$ | 3.7±0.009$\times 10^{-2}$ |
| Half-life($t_{1/2}$) | 18.5 min | 17.3min | 18.2min |

#### 3.4.2 H-5A and H-38A mutations reduce hydroxylamine (HA) induced N-cleavage

HA is an alpha nucleophile that intercepts between amide-ester equilibrium and induces N-cleavage. It enhances the nucleophilicity of residues that leads to N-cleavage (84, 85). A pseudo-first-order fit for the HA-induced reaction time course with the equation, Y=Y0 +(Plateau-Y0) *(1-exp(-K*x)) was obtained (Figure 5). H-5A and H-38A mutants exhibited about 1.4-fold and 1.5-fold reduction (p=0.0025; one-way ANOVA) in % of N-cleavage, respectively, relative to *Mtu* FL-SufB although the K and t1/2 values were comparable [Figure 5C, Table 2 (ii)]. This effect on HA-induced N-cleavage reaction was similar to what is observed in presence of TCEP for the H-5A and H-38A mutations.

#### 3.4.3 H-5A and H-38A mutations do not alter DTT induced N-cleavage

Similar to HA, DTT is also a nucleophile that enhances the nucleophilicity of residues and facilitates N-terminal cleavage reaction (83, 86). N-cleavage efficiency was not noticeably different between H-5A and H-38A mutants and *Mtu*-FL-SufB in presence of DTT (Figure S3 and Table S4). DTT interacts directly with the residues that may increase the flexibility of the active site structure (87), which possibly counteracts the effect of H-5A and H-38A mutations on N-terminal cleavage. The hypothesized role of the histidines at -5 and -38 positions in the N-extein region in direct or indirect Cys1 activation was probed further by MD simulations.

### 3.5 Protein identification

The identity of splicing and cleavage products (P, CC, LE, and NE) for the N-terminal 6X (His) tagged proteins was affirmed via western blot using anti-His antibodies (Figure 3B, 5B, and S3B). SI (C1A/N359A) double mutant SufB was used as a control that yielded just unspliced precursor protein (P) [Figure 3B (iv)]. One additional protein band was noticed above LE and below CC for *Mtu* FL-SufB, H-5A, and H-38A possibly due to a splicing intermediate as an off-pathway product. We also observed a product just below the P band for *Mtu* FL-SufB, SI, H-5A, and H-38A (Figure3B, 5B, and S3B). However, this protein band was missing in cells expressing SufB intein or empty vector pACYC Duet-1 [Figure 3B (iv)]. This indicated that this band corresponded to a SufB precursor degradation product in the cells over-expressing active and inactive *Mtu* FL-SufB precursor.

To further clarify the identity of these different splicing and cleavage products, MALDI-TOF/TOF mass spectrometry (MS) was performed (Table S3) on protein bands cut from the SDS PAGE gel. When the acquired MS and MS/MS spectra were searched against the taxonomy Mycobacterium tuberculosis from NCBInr (88) database, all the splicing products were identified as SufB protein of *Mycobacterium tuberculosis* complex with a statistically significant score (Table S3). A further check against a customized database containing probable splicing and cleavage products of SufB protein of *Mycobacterium tuberculosis* with same parameters for the individual protein band spectra resulted in the identification of matching protein splicing products. Individual protein band identifications with protein score, percent coverage, theoretical molecular weight, and isoelectric pH value are shown in Table S3. Significance is measured from the expectancy-value (with a p-value ≤0.05).

Protein fragments of *Mtu* FL-SufB were also found to be within the expected mass range as per retention time (RT) by HPLC (Figure S1).

### 3.6 Proposed mechanism for the cleavage of N-terminal intein∼extein peptide bond

The *in vitro* experimental results suggest that the mechanism of *Mtu* SufB intein cleavage at the N-terminal cleavage site between Gly252 (G-1) and Cys253 (C1) is influenced by two conserved histidines at -5 (248, full-length protein) and -38 (215, full-length protein) positions in the N-extein sequence of the *Mtu* FL-SufB protein. To further probe the basis of this influence, we performed MD simulations on a 3-dimensional (3D) model of the *Mtu* FL-SufB precursor protein (Figure. 6A). This also aided identification of possible critical residues at the SufB intein active site (N- and C-terminal splice junctions) (Figure. 6B and Figure. S7), given their proximity to catalytic residues Cys253 (C1), Cys612 (C+1), and Asn611 (N359).

**Figure 6.**
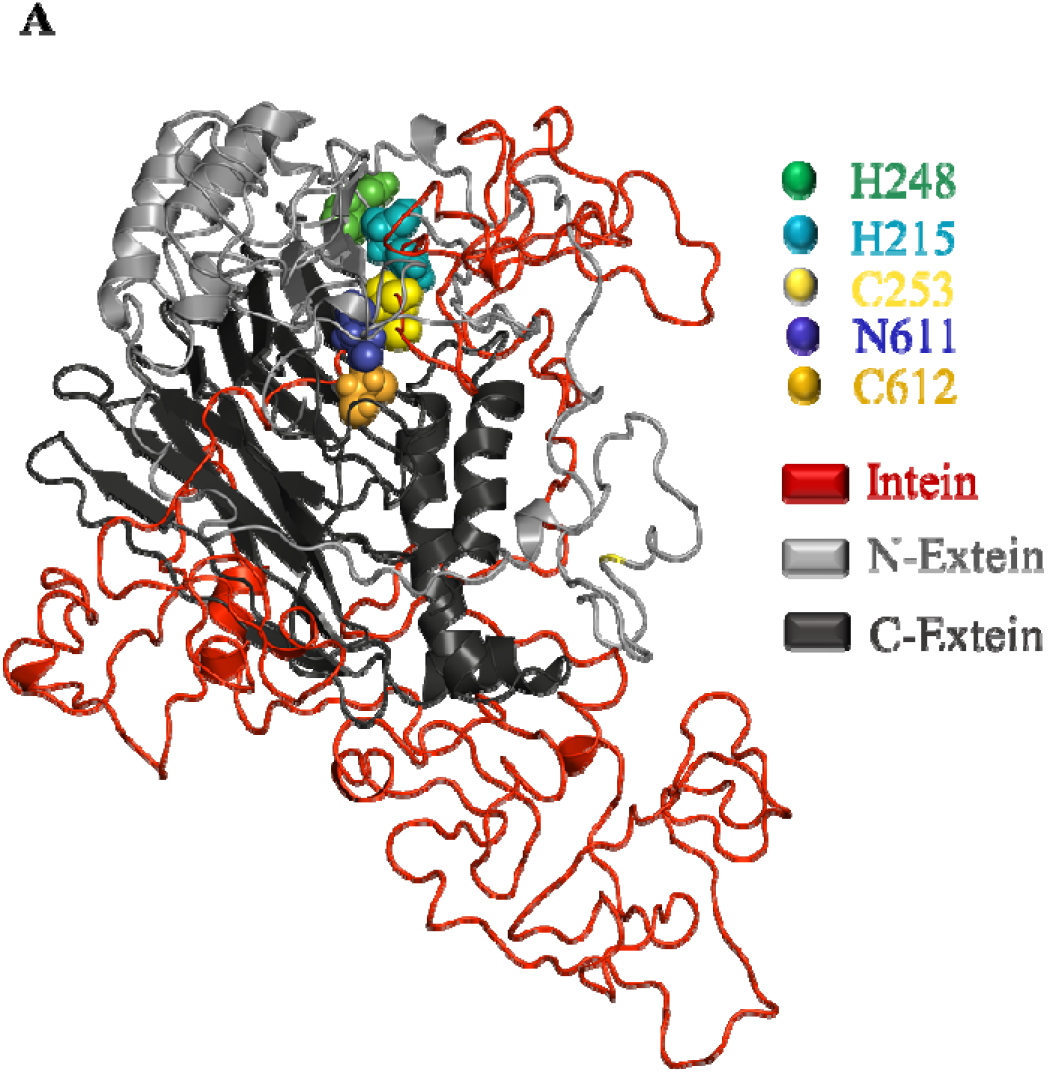
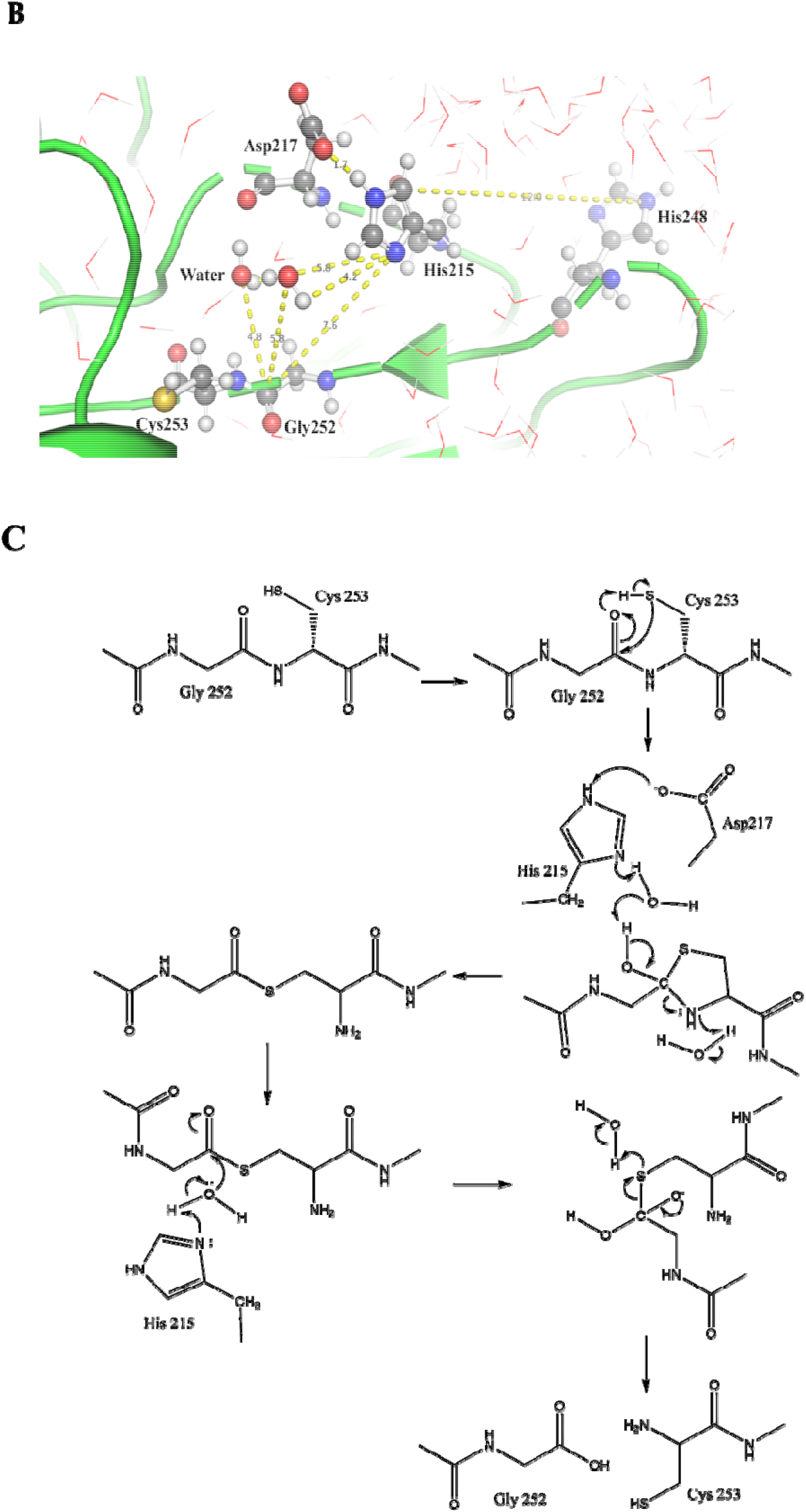
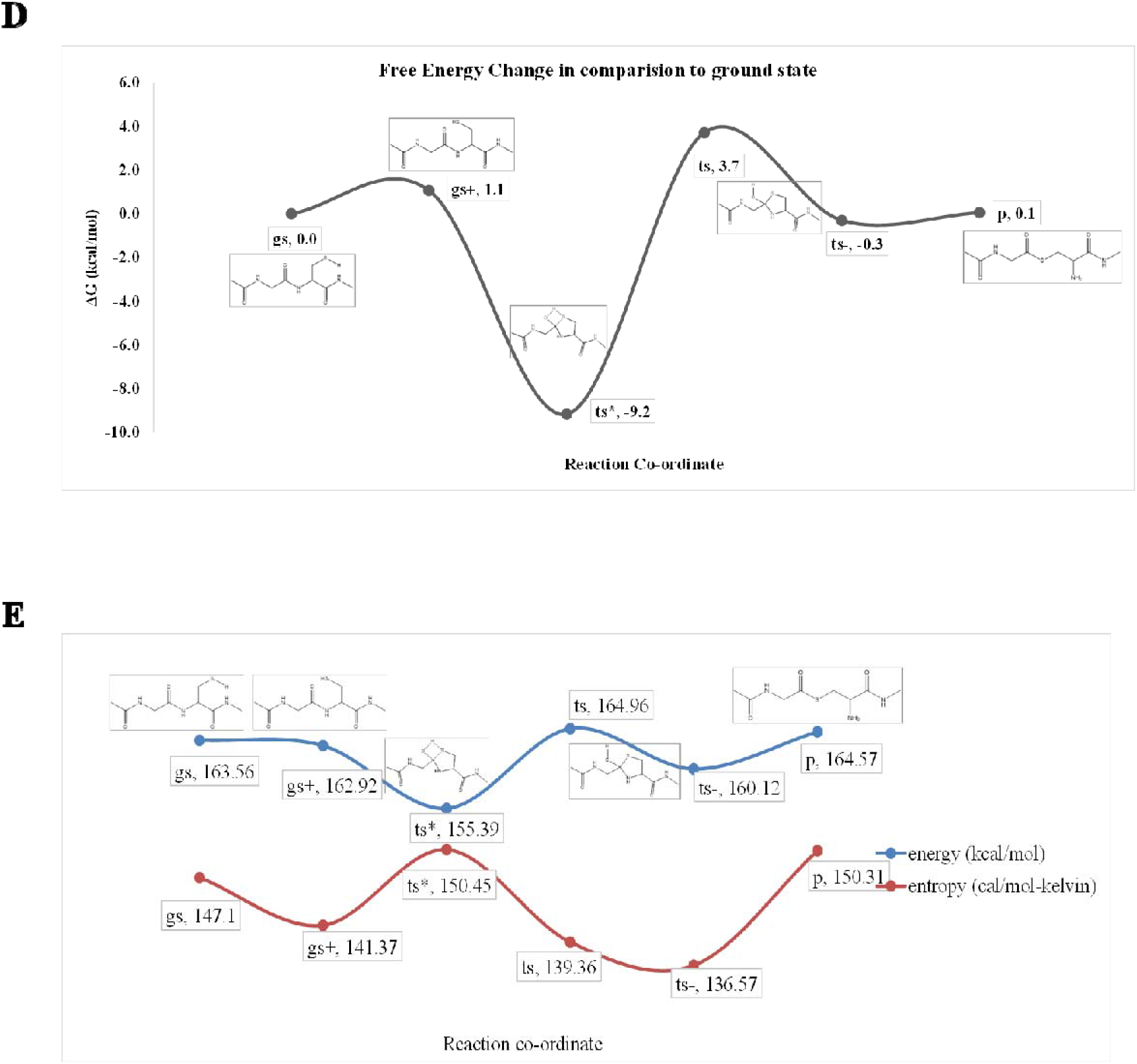
Structural model and proposed mechanism for N-cleavage reaction in *Mtu* FL-SufB precursor. (A) The 3D model of *Mtu* FL-SufB precursor obtained in explicit solvent. Different domains and conserved catalytic residues are color coded as shown in the legend, (B) Spatial arrangement of the N-terminal active site residues; Cys253 (C1) and Gly252 (G-1) along with His248 (H-5), His215 (H-38) and Asp217 (D-36) at the N-terminal cleavage site, (C) Proposed mechanism for N-terminal cleavage reaction at intein∼extein junction, (D) Free energy change, and (E) Energy and entropy of the system during QM calculation of different reaction states of N-cleavage junction from peptide bond to thio-ester bond of *Mtu* FL-SufB precursor protein. gs = ground state, gs+ = conformational transition state, ts*= transition stat of proton transfer, ts = tetrahedral intermediate, ts = transition state of proton transfer, p = product.

The composite 3D model of *Mtu* FL-SufB precursor was built based on percent sequence identity with analogous sequences from the SufBCD complex of *Escherichia coli* (40% identity, PDB ID: 5AWF, chain A), PI-SceI; a homing endonuclease with protein splicing activity (45% identity, PDB ID: 1VDE, chain A) and intein homing endonuclease II (22% identity, PDB ID: 2CW7, chain A). This model of *Mtu* FL-SufB lacked secondary structure in the intein region due to low sequence identity to the template, with nearly 13% of amino acids are found to be present in the disallowed region in the Ramachandran plot (Figure S5A). To optimize this structure further, we carried out molecular dynamics (MD) simulations of the model in explicit solvent for a duration of 100ns. Based on parameters such as solvent accessible surface area (SASA) (Figure. S4B) and root mean square deviation (RMSD), the structure was found to be equilibrated after 80ns (Figure. S4A). The equilibrated *Mtu* FL-SufB structure was found to have only ∼1% amino acids in the disallowed region of the Ramachandran plot (Figure. S5B). We found that the longer side chain amino acids in the disallowed region of the Ramachandran plot were mostly solvent-exposed. The ProSAweb score of the model before and after the 100 ns MD simulations is shown in Figure. S5C and D.

The simulations were also analyzed for the role of specific residues, especially His248 (H-5), which is near the N- and C-cleavage sites. The comparison between 20 ns duration simulations for the *Mtu* FL-SufB precursor and H-5A mutant for RMSD and Root Mean Square Fluctuation (RMSF) parameters is shown in Figure. S6A and B. It was observed that His248 (H-5) attracts His215 (H-38) towards the active site Gly252 (Gly-1), which could facilitate the *Mtu* SufB N-cleavage reaction. Although His-5 is closer to the active site compared to His-38) in the primary sequence, His-38 is closer to the active site in the 3D model. Specifically, the shortest distance between His-38 and Gly-1 was observed to be 6.8 Å, which is similar to distances observed between catalytic residues in other inteins (12, 39).

Similarly, His610 (His358), Asn 611 (His359), and Cys612 (Cys+1) were located near the C-cleavage site in the *Mtu* FL-SufB MD simulations (Figure S7). The distance between Asn611 (N359) and Cys612 (C+1) was found to be 5.5 ± 0.1 Å and the distance between Asn611 (N359) and His610 (H358) was 4.3 ± 0.3 Å (Table S5). This proximity suggests that the C-cleavage reaction may be facilitated by interactions between these three amino acids, but further analysis of this reaction step is outside the scope of the present study.

Coordination between N-terminal intein∼extein splice junction residues Gly252 (Gly-1), Cys253 (Cys1), and His215 (His-38) and His248 (His-5) likely propels *Mtu* SufB N-terminal cleavage. Quantum mechanical (QM) calculations were used to obtain energetics of this N-cleavage reaction as done before for intein cleavage of the *Mtu* RecA intein (23). After the optimized structures of the reactants and the products for each step were obtained, they were used as the initial and final states respectively. Frequency analyses helped identify the stationary points of the intermediates (only real frequencies) and the saddle points of the transition states (one imaginary frequency). The reaction energy barrier was obtained from the energy difference between the transition state and reactant state with the zero-point correction. A postulated schematic for the cleavage reaction, analogous to the canonical intein splicing mechanism (89), is shown in Figure 6C. All stationary states were confirmed by means of vibrational frequency analysis, and the Gibbs free energy at physiological temperature was calculated for all stationary points, including the zero-point energy, the entropy, and the thermal energy (all at 298.15K) at the B3LYP/6-31G (d) (Figure 6D, 6E).

## 4. DISCUSSION

### 4.1. Mtu SufB likely follows a canonical intein splicing mechanism

Being an essential component of the mycobacterial SUF system, the SufB protein is well conserved in mycobacteria, bacteria, and archaea. Intein-carrying SufB proteins are found in different mycobacterial species, although the intein insertion points vary in some species (Figure 2) (54). Cladogram analysis suggest some divergence in *M. leprae*, *M. lepromatosis*, *M. xenopi*, *M. triplex* and Ferroplasma in the extein sequences and intein insertion sites, which suggests intein-extein co-evolution and independent intein transfer in different species and kingdoms. Domain analysis of the *Mtu* FL-SufB precursor (846 aa) clearly demarcates the intein and extein structural regions (Figure 1, Table S2), with the *Mtu* SufB intein (359 residues) containing an intact endonuclease domain. Catalytic residues critical for classic intein splicing pathways like Cys1 (Block A), Cys+1 (Block G), penultimate His (Block G), and terminal Asn (Block G) are conserved in different mycobacterial species including *Mtu*. The *Mtu* SufB intein also contains the conserved Block B His67 that is known to catalyze the first N-S acyl shift by destabilizing the scissile peptide bond. In vitro refolding of *Mtu* SufB precursor gives splicing and cleavage products that also concur with the roles of these conserved residues. All these observations suggest that *Mtu* FL-SufB undergoes a canonical (classic) cis-splicing autoprocessing mechanism to form the functional ligated extein protein (Figure 1).

### 4.2. A distinct N-terminal cleavage mechanism regulated by conserved H-5 and H-38

Sequence analysis detected two highly conserved His residues in the N-extein sequence of *Mtu* SufB protein; His-38 and His-5 located in all SufB proteins where SUF constitutes the exclusive pathway for [Fe-S] cluster generation irrespective of genus and kingdoms (Figure 2A, B). Catalytic residues such as Cys1, Cys+1, and terminal Asn directly participate in intein splicing by promoting sequential nucleophilic displacement reactions or by rearrangement of bonds near splice junctions. Non-catalytic residues assist indirectly via activation of active site residues and stabilization of various intermediate structure(s). Conserved His residues within intein sequence are known to play important roles during protein splicing as well. Block B His accelerates N-S acyl shift and cleavage of N-terminal intein∼extein peptide bond whereas F- and G-Block His are crucial in the coordination of terminal Asn cyclization and cleavage of C-terminal splice site.

Splicing assays and kinetic analysis on *Mtu* FL-SufB precursor and mutant proteins suggests a mechanism where His-5 and His-38 coordinate to facilitate N-cleavage reaction via activation of catalytic Cys1 [Figure 3, 4, and 5]. Both H-5A and H-38A mutants exhibit a sluggish splicing reaction relative to FL-SufB protein under optimum experimental conditions (Table 1). N-cleavage reaction kinetic analysis shows that H-5A and H-38A mutations reduce N-cleavage efficiency in presence of TCEP and HA (Table 2 (i) and (ii), Figure 4 and 5), but DTT-induced thiolysis is unaffected (Figure S3). This may be explained by DTT being a stronger nucleophile that exhibits a more efficient thiolysis and di-sulfide reduction. H-5 and H-38 therefore seem to assist *Mtu* SufB N-cleavage in conjunction with other active site residues.

To further ascertain the roles of these conserved N-extein histidines, a 3D model of the *Mtu* FL-SufB precursor was built by homology modeling. Explicit solvent MD simulations were performed to optimize and equilibrate the model. H-38 was observed to be localized closer to the active site in the optimized model, which suggests that H-5 and H-38 act in a concerted manner to facilitate *Mtu* SufB N-cleavage. Other active site residues such as Gly-1 and Cys1 also seem to be involved. QM/MM calculations using equilibrated structures from the MD simulations provide a quantitative characterization of the energetics of the N-cleavage reaction.

Taken together, these results support the following postulated mechanism for *Mtu* SufB N-cleavage reaction, which is shown as a schematic in Figure 6C. The polarized thiol group of Cys1 approaches the peptidyl C=O, with this transition state being enthalpically favored by an interaction between the thiol H and the peptidyl O atom. The electron density decreases in the C, S, and H atoms while increasing in the O atom. With the imaginary vibrational frequency displacement vector of C=O being towards the thiol, the C and S atoms come closer to form a C-S bond. The proton migration from S to O is highly favored energetically and entropically with a free energy change of -9.2 kcal/mol, which agrees with earlier results (90, 91).The sp2 hybridized peptide carbon becomes sp3 hybridized forming a tetrahedral transition state with a 2-hydroxy thiazolidine ring. This annihilates the peptide resonance to increase the free energy by +3.7 kcal/mol. To regain the sp2 hybridization and resonance stabilization, the 2-hydroxy group expels proton to the environment. In a tetrahedral state, S and N atoms have a Mulliken charge of +0.05 and -0.55. The N atom of the thiazolidine ring accepts a proton from the environment become neutral and forms a thioester intermediate (90, 92). The conversion of a peptide bond to a thioester bond is energetically equivalent at the start as well as the end of the reaction which suggests this process is entropy-driven. The thioester could be hydrolyzed by the imidazole side chain of His-38 mediated by water. Such base catalyzed thioester hydrolysis in an aqueous environment is well known (93). The least distance between His-38 and His-5 in MD simulations is about 6.8 Å suggesting an interaction between these two residues to aid catalysis (12, 19).

Asp-36 could exchange a proton with the His-38. The distance fluctuations between His-38 and His-5 in the MD simulations suggest that the solvent-exposed His-5 could attract His-38 towards the N-cleavage site through water-mediated interactions. His-5 could then facilitate thioester hydrolysis by the attacking species (i.e., hydroxide ion) being generated by His-38.

### 4.3. Biological significance of conserved histidines in the N-extein sequence of Mtu SufB

[Fe-S] cluster-bearing proteins have important physiological roles in electron transfer, redox regulation, metabolic pathways, cellular responses to external stimuli, and as regulators of gene expression (45). The role of the [Fe-S] cluster is closely associated with the functionality of their bound protein framework. In mycobacteria, the SUF system is the sole pathway for [Fe-S] cluster assembly and repair, especially in response to oxidative stress and iron limiting conditions inside macrophages (46, 52). During such an event, the intracellular Fe supply is either from siderophore chelation or via the metabolism of [Fe-S] clusters associated with specific proteins (45). Thus, iron homeostasis plays a major role in mycobacterial survival and virulence. SufB is a [Fe-S] cluster scaffold protein and a vital component of the functional SUF system, and indirectly promotes mycobacterial persistence under stress(52). Further, SufB has been implicated in mycobacterial iron metabolism(45).

Earlier studies have specified the bonding of Fe in [Fe-S] clusters to mostly cysteines from the protein backbone although there is increasing evidence for other ligands such as histidine, aspartate, arginine, threonine, and tyrosine. The most common alternative ligand for [Fe-S] cluster coordination is histidine that is highly conserved with a role in redox tuning and proton-coupled electron transfer (94). His433(SufB) and His360(SufD) are identified as key protein-ligands for the *de novo* [Fe-S] cluster assembly in *E. coli* (51). It has been shown that these non-cysteine ligands can influence the stability and reactivity of [Fe-S] clusters.

In addition to their possible role in initiating N-cleavage reaction via catalytic Cys1 activation, the conserved His-5 and His-38 residues in the N-extein sequence of *Mtu* SufB may play a role in precursor stabilization. Apart from Cys, His is also an important ligand for metal coordination (51, 94), and His-5 and His-38 could coordinate to Fe+2/ Fe+3 during [Fe-S] cluster biogenesis. Confirmation of SufB precursor stabilization by coordination of these His residues to [Fe-S] cluster or to Fe^+2^/ Fe^+3^ would need further studies.

SUF is an exclusive system for biogenesis of Fe-S clusters in many pathogenic organisms such as *Staphylococcus aureus*, *Mycobacterium tuberculosis*, *Plasmodium sp.* and *Toxoplasma*, making it an attractive drug target. For instance, D-Cycloserine is a clinical second-line drug currently used against *M. tuberculosis* and inhibits SufS. SUF is the target system for a polycyclic molecule 882 that has direct interaction with SufC in *S. aureus*(80). The current work may therefore aid development of novel anti-TB drugs targeting SufB function and stability.

## Data Availability

All the data are available in the manuscript and supplemental data.

The accession code for the *Mycobacterium tuberculosis* SufB protein used in this study is *P9WFP7 (UniProtKB/Swiss-Prot) and* WP_003407484.1, GI: 397673309 (NCBI protein database).

Confirmation of the identity of different splicing and cleavage products of Mtu FL-SufB protein by mass spectrometric analysis: The data has been deposited to the ProteomeXchange Consortium via PRIDE partner repository with data set identifier PXD015199. The details of the submission are given below.

**Project Name**: Identification of full length, splicing and cleavage products of SufB protein of SUF-complex of *Mycobacterium tuberculosis*.

Project accession: PXD015199

Reviewer account details:

**Username**

**Password**: bRxRcbvP

The homology model for the full length Mtu SufB protein has been submitted to Model Archive [Project: ma-x807d]:

Full length *Mtu* SufB protein of SUF complex of *Mycobacterium tuberculosis*; [https://www.modelarchive.org/doi/10.5452/ma-x807d] with the access code: 6pmXRNkvwR.

## Acknowledgements

Our sincere thanks to Prof. Marlene Belfort, Department of Biological Sciences and RNA Institute, University at Albany, Albany, NY, USA, for her valuable comments on the manuscript. The plasmids used in this study are borrowed from Prof. Marlene Belfort’s laboratory, University at Albany, Albany, NY, USA. These were engineered by Dr. Sasmita Nayak during her PhD work under the guidance of Prof. Marlene Belfort. We also thank Mr. Sourya Subhra Nasker for providing critical comments on the manuscript. We are thankful to Mr. Rajendra Reddy for technical support at Central Proteomics Facility, Institute of Life sciences, and Bhubaneswar. We also thank Mr. A. Ananda Raman for his timely help in HPC (High Performance computing cluster) to perform computational work at National Institute of Science Education and Research (NISER) Bhubaneswar. Our sincere gratitude to Mrs. Anuradha Das for her help in conducting the HPLC experiment at School of Chemical Sciences, NISER Bhubaneswar and to Dr. Rahul Modak for his help in conducting FPLC at School of Biotechnology, KIIT Deemed to be University.

## Author contributions

**Dr. Sunita Panda**: Methodology; Validation; Formal analysis; Investigation; Resources; Data curation; Writing - original draft; Visualization; Funding acquisition. **Ananya Nanda:** Methodology; Validation; Formal analysis; Investigation; Resources; Data curation; Writing - original draft; Writing- Reviewing and Editing; Visualization. **Nilanjan Sahu:** Methodology; Software; Validation; Formal analysis; Writing - original draft. **Deepak Kumar Ojha:** Validation; Formal analysis; Investigation; Resources. **Dr. Biswaranjan Pradhan:** Formal analysis; Resources. **Anjali Rai**: Resources. **Dr. Amol R. Suryawanshi:** Methodology; Validation. **Dr. Nilesh Banavali**: Methodology; Software; Validation; Writing- Reviewing and Editing; Visualization. **Dr. Sasmita Nayak:** Conceptualization; Methodology; Validation; Investigation; Data curation; Writing - original draft; Writing- Reviewing and Editing; Visualization; Supervision; Funding acquisition; Project administration.

## Abbreviations

Mtu: Mycobacterium tuberculosis
WT: Wild type
SI: Splicing inactive
DM: Mtu SufB Double mutant
IPTG: Isopropyl β-d-1-thiogalactopyranoside
TCEP: Tris (2-carboxyethyl) phosphine
HA: Hydroxylamine
DTT: Dithiothreitol
HRP: Horseradish peroxidase
ECL: Enhanced chemiluminescence
TBST: Tris-buffered saline, 0.1% tween 20
BSA: Bovine serum albumin
ß-gal: Beta galactosidase
ABC: Ammonium bicarbonate
ACN: Acetonitrile
TFA: Tri-fluoro acetic acid
HPLC: High pressure (or performance) liquid chromatography
FPLC: First protein liquid chromatography
MALDI-TOF/TOF: Matrix assisted laser desorption ionization time-of-flight
SDS: Sodium dodecyl sulphate
PAGE: Polyacrylamide gel electrophoresis
RMSD: Root mean square deviation
SASA: Solvent accessible surface area
RMSF: Root mean square fluctuation
PME: Particle-mesh Ewald
HPCC: High performance computing cluster
MD Simulations: Molecular dynamic simulations

