## Supplemental Files for "SufB intein splicing in *Mycobacterium tuberculosis* is influenced by two remote conserved N-extein Histidines"

**Table of Contents: Page No.**

**Table S1**: List of primers for 6X(His)- tagged *Mtu* full-length (FL) *sufB* gene and	S3

phosphorylated primers for mutagenesis PCR.

**Table S2:** Amino acid sequences of different structural domains of *Mycobacterium*	S4	*tuberculosis (Mtu)* full-length SufB precursor protein.

**Table S3**: MALDI-TOF MS protein identification using *Mycobacterium tuberculosis*	S5

complex, from NCBI nr protein database.

Determination of relative molecular weight of different splicing and cleavage products	S6

of *Mtu* full-length (FL) SufB via HPLC analysis.

**Figure S1:** Determination of relative Molecular weight of *Mtu* FL-SufB	S6

fragments via HPLC.

Analysis of splicing inactive (SI) double mutant (C1A/N359A) *Mtu* SufB expression over a period of 24hours	S7

**Figure S2:** Effect of C1A/N359A double mutation on *Mtu* FL-SufB splicing and	S7

N-cleavage reaction at 20^0^C.

**Figure S3**: Comparison of DTT induced N-terminal cleavage reactions in *Mtu* FL-SufB precursor, H-5A and H-38A mutant SufB proteins.	S8

**Table S4:** Analysis of different kinetic parameters for DTT induced N-cleavage	S9

reaction in FL-SufB, H-5A and H-38A SufB mutants.

**Figure S4:** Analyses on Root Mean Square Deviation (RMSD) and Solvent Accessible	S9

Surface Area (SASA) of *Mtu* FL-SufB protein.

**Figure S5:** Model validation of proposed *Mtu* FL-SufB precursor structure. S10

**Figure S6**: Analysis on comparison of RMSD of *Mtu* FL-SufB and H-5A SufB	S10

mutant and RMSF of FL-SufB and H-5A SufB mutant.

**Figure S7:** Structural model showing spatial arrangement of active site	S11

residues at C-terminal cleavage junction.

**Table S5:** Molecular Dynamics analysis of *Mtu* FL-SufB showing average	S11

distance between center of mass of active site residues at C-terminal intein~extein junction.

-----------------------------------------------------------------------------------------------------------------

**Table S1. *List of primers for 6X(His)- tagged Mtu FL-sufB gene and phosphorylated primers for mutagenesis PCR. P~ denotes 5’ phosphate.***

| ***Primer Name*** | ***Primer sequence*** |
| --- | --- |
| ***Mtu _FL_SufB_F*** | ***GCATGCGAATTCGATGACACTCACCCCAGAGGCC*** |
| ***Mtu _FL_SufB_R*** | ***GTCATGCAAGCTTTCATCCGACCGCGCCCTCCATCTG*** |
| ***Mtu _FL_SufB_His_F*** | ***GCATGCGAATTCGGATGACACTCACCCCAGAGGCC*** |
| ***Mtu _FL_ SufB _C1A_ F*** | ***P~GCGCTGCCCGCCGGCGAGCTCATCACG*** |
| ***Mtu _FL_ SufB _C1A_R*** | ***P~GCCCTCTACGTAGTGCACGTAAGAGCC*** |
| ***Mtu _FL_ SufB _N359A_F*** | ***P~TGCACCGCACCGATCTACAAATCGGATTC*** |
| ***Mtu _FL_ SufB _N359A_R*** | ***P~CGCGTGCACGGCGAACCCGTAGGCGAG*** |
| ***Mtu _FL_ SufB _C+1A_F*** | ***P~GCGACCGCACCGATCTACAAATCGGATTCATTG*** |
| ***Mtu _FL_ SufB _C+1A_R*** | ***P~GTTGTGCACGGCGAACCCGTAGGC*** |
| ***Mtu _FL_ SufB _H-5A_F*** | ***P~GCGTACGTAGAGGGCTGCCTGCCC*** |
| ***Mtu _FL_ SufB _H-5A_R*** | ***P~CACGTAAGAGCCCTCATCGGCG*** |
| ***Mtu _FL_ SufB _H-38A_F*** | ***P~GCG GTC GAC ATT CCG CTG CA*** |
| ***Mtu _FL_ SufB _H-38A_R*** | ***P~AAC ACC GGG CGG GAC GTA AAT*** |

**Table S2. *Amino acid sequences of different structural domains of Mycobacterium tuberculosis(Mtu) full-length (FL) SufB precursor protein:***

***Mtu SufB N-extein:*** MTLTPEASKSVAQPPTQAPLTQEEAIASLGRYGYGWADSDVAGANAQRGLSEA VVRDISAKKNEPDWMLQSRLKALRIFDRKPIPKWGSNLDGIDFDNIKYFVRSTEK QAASWDDLPEDIRNTYDRLGIPEAEKQRLVAGVAAQYESEVVYHQIREDLEAQG VIFLDTDTGLREHPDIFKEYFGTVIPAGDNKFSALNTAVWSGGSFIYVPPGVHVDI PLQAYFRINTENMGQFERTLIIADEGSYVHYVEG

***Mtu SufB intein, N-terminal domain:***

CLPAGELITTADGDLRPIESIRVGDFVTGHDGRPHRVTAVQVRDLDGELFTFTPM SPANAFSVTAEHPLLAIPRDEVRVMRKERNGWKAEVNSTKLRSAEPRWIAAKDV AEGDFLIYP

***Mtu SufB intein, Homing*** ***endonuclease domain:***

KPKPIPHRTVLPLEFARLAGYYLAEGHACLTNGCESLIFSFHSDEFEYVEDVRQAC KSLYEKSGSVLIEEHKHSARVTVYTKAGYAAMRDNVGIGSSNKKLSDLLMRQD ETFLRELVDAYVNGDGNVTRRNGAVWKRVHTTSRLWAFQLQSILARLGHYATV ELRRPGGPGVIMGRNVVRKDIYQVQWTEGGRGPKQARDCGD

***Mtu SufB intein, C-terminal domain:*** YFAVPIKKRAVREAHEPVYNLDVENPDSYLAYGFAVHN

***Mtu SufB C-extein:*** CTAPIYKSDSLHSAVVEIIVKPHARVRYTTIQNWSNNVYNLVTKRARAEAGATM EWIDGNIGSKVTMKYPAVWMTGEHAKGEVLSVAFAGEDQHQDTGAKMLHLAP NTSSNIVSKSVARGGGRTSYRGLVQVNKGAHGSRSSVKCDALLVDTVSRSDTYP YVDIREDDVTMGHEATVSKVSENQLFYLMSRGLTEDEAMAMVVRGFVEPIAKE LPMEYALELNRLIELQMEGAVG

***MALDI-TOF MS protein identification using Mycobacterium tuberculosis complex, from NCBI nr protein database:***

MALDI-TOF/TOF mass spectrometry was performed for confirmation of different splicing and cleavage products generated from *Mtu* FL-SufB precursor protein (Table S3). The details of procedures and analyses are provided in the main text (Section 2.5 and Section 4.7)

**Table S3. *MALDI TOF/TOF MS identification of Mtu FL-SufB protein, splicing and cleavage products from protein pilot database****.* Previously all the protein spots were identified as full-length (FL) SufB from *Mycobacterium tuberculosis complex.* From the identified protein, individual domains or splicing products were selected and the respective FASTA files were submitted with accession number to protein pilot database. Then respective proteins and peptides were identified by PMF search with best protein score, best protein mass and with best protein description.

| Protein Spot | Best Protein Accession | Best protein mass (Dalton) | Best Protein Score | Best protein Description |
| --- | --- | --- | --- | --- |
| A15 (Intein) | Wt_I_WP_003407484.1 | 40546 | 127 | Fe-S Cluster assembly protein SufB  [*Mycobacterium tuberculosis*] |
| A16 (Precursor) | Wt_P_WP_003407484.1 | 94852 | 633 | Fe-S Cluster assembly protein SufB  [*Mycobacterium tuberculosis*] |
| A21 (N-Cleavage product) | Wt_NC_WP_003407484.1 | 65258 | 355 | Fe-S Cluster assembly protein SufB  [*Mycobacterium tuberculosis*] |
| A22 (Ligated Extein) | Wt_LE_WP_003407484.1 | 53805 | 334 | Fe-S Cluster assembly protein SufB  [*Mycobacterium tuberculosis*] |
| B2 (N-extein) | Wt_NE_WP_003407484.1 | 27957 | 263 | Fe-S Cluster assembly protein SufB  [*Mycobacterium tuberculosis*] |

***Determination of relative molecular weight of different splicing and cleavage products of Mtu* FL*-*SufB *via HPLC analysis:***

The reaction volume containing the refolded protein fragments of *Mtu* FL-SufB was subjected to HPLC analysis. The retention time vs protein intensity measured at 280nm was noted. For reference, we have used Precision plus protein ladder (Biorad1610374). The retention time Vs molecular weight of the known protein standards was measured and plotted to make a standard curve. The unknown protein peaks from the test samples was compared with the standard curve to find out the relative molecular weights. The molecular weights of the expected fragments from the MALDI-TOF/TOF MS (Table S3) data were compared with the standard curve and molecular weights of the unknown peaks were determined.

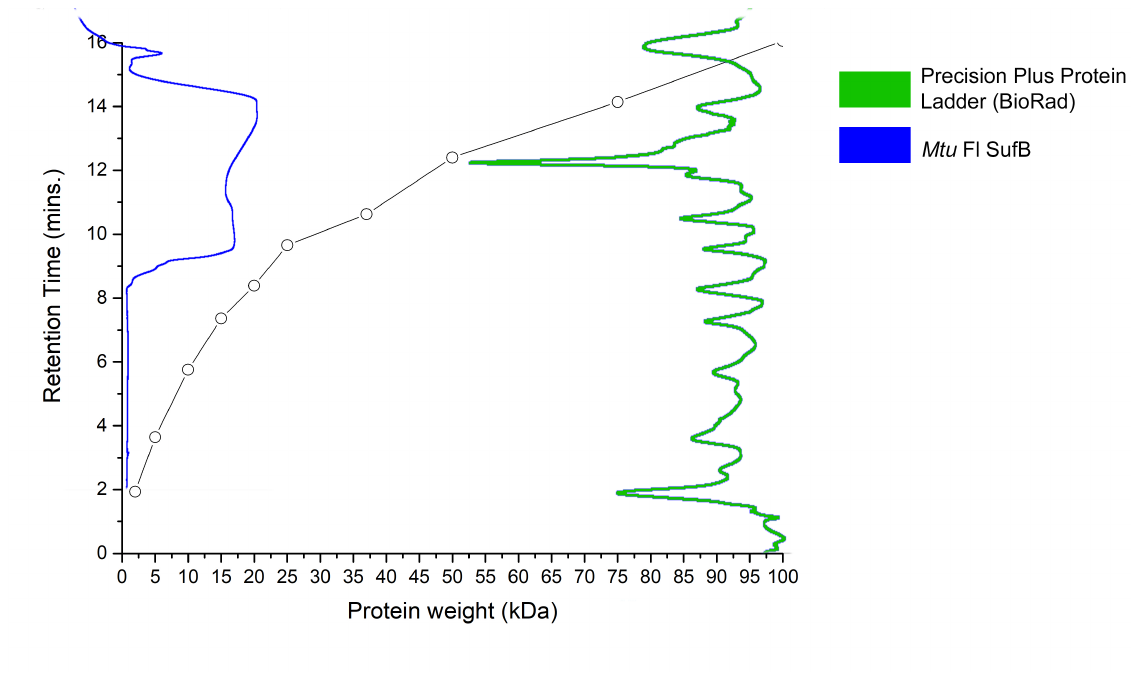

**Figure S1. Determination of relative molecular weight of *Mtu* FL-SufB fragments.** Molecular weight of protein vs retention time of Precision plus Protein Ladder (BioRad) was plotted as a standard curve for comparison. In-plot, to the right (Green) is the chromatogram of aforesaid protein ladder, to the left, plot depicting different fractions of *Mtu* FL-SufB (Blue).

***Analysis of splicing inactive (SI) double mutant (C1A/N359A) SufB expression over a period of 24hours:***

The splicing inactive (SI) SufB double mutant, with both C1 and N359 mutated to alanine (C1A/N359A), which is expected to abolish intein splicing completely, was used as a negative control for splicing. Transformants expressing empty vector pACYC Duet-1 were used a negative control for protein expression. Products from *in-vitro* refolding assay for a period of 24hours were resolved through 4~10% gradient SDS PAGE as explained in the main text (Materials and Methods). As expected, none of the post-reaction products were seen in the case of SI double mutant (Figure S2A) and cells expressing empty vector pACYC Duet-1 (Figure S2B).

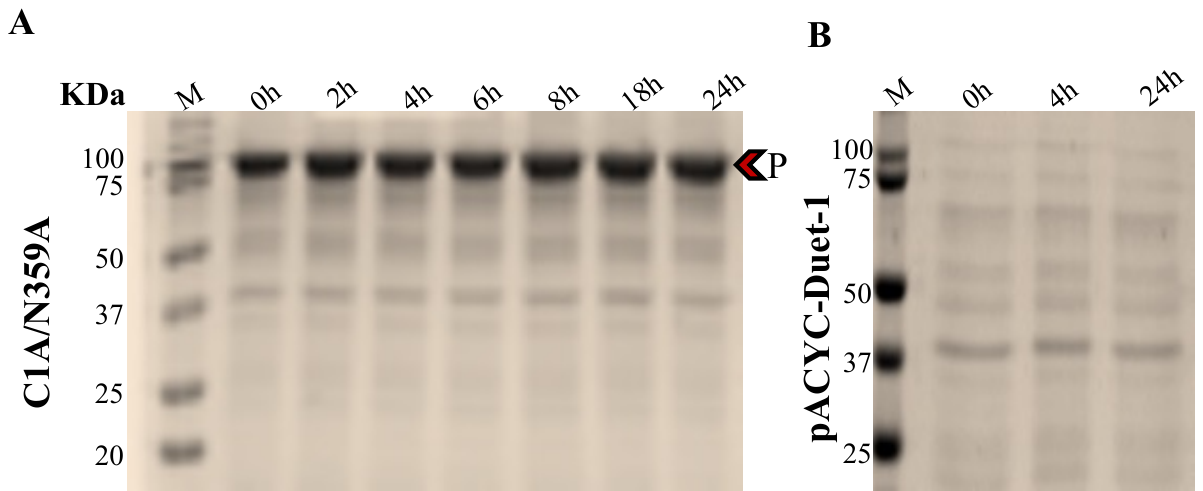

**Figure S2. Effect of C1A/N359A double mutation on *Mtu* SufB splicing and N-cleavage reaction at 20˚C.** Products from *in-vitro* refolding assay for a period of 24hours were resolved through 4~10% gradient SDS PAGE. (A) SI (Splicing inactive C1A/N359A SufB double mutant) and (B) pACYC Duet-1 empty vector were used as negative controls for the above study. P: SI SufB precursor protein.

***Kinetic assay of DTT induced N-cleavage reactions in Mtu FL -SufB, H-5A and H-38A SufB mutant proteins:***

For better understanding of the reactions steps for our test proteins, we performed an *in-vitro* refolding and N-cleavage assay of DTT induced reactions. DTT being a thiolate has dual properties; it can act both as nucleophile and reducing agent. Since the optimum temperature for FL-SufB splicing (section 2.3) was found to be 20°C, N-cleavage reactions for the above proteins were also conducted at the same temperature. The details of the procedures and data analyses for the above study are provided in the main text (Materials and Methods).

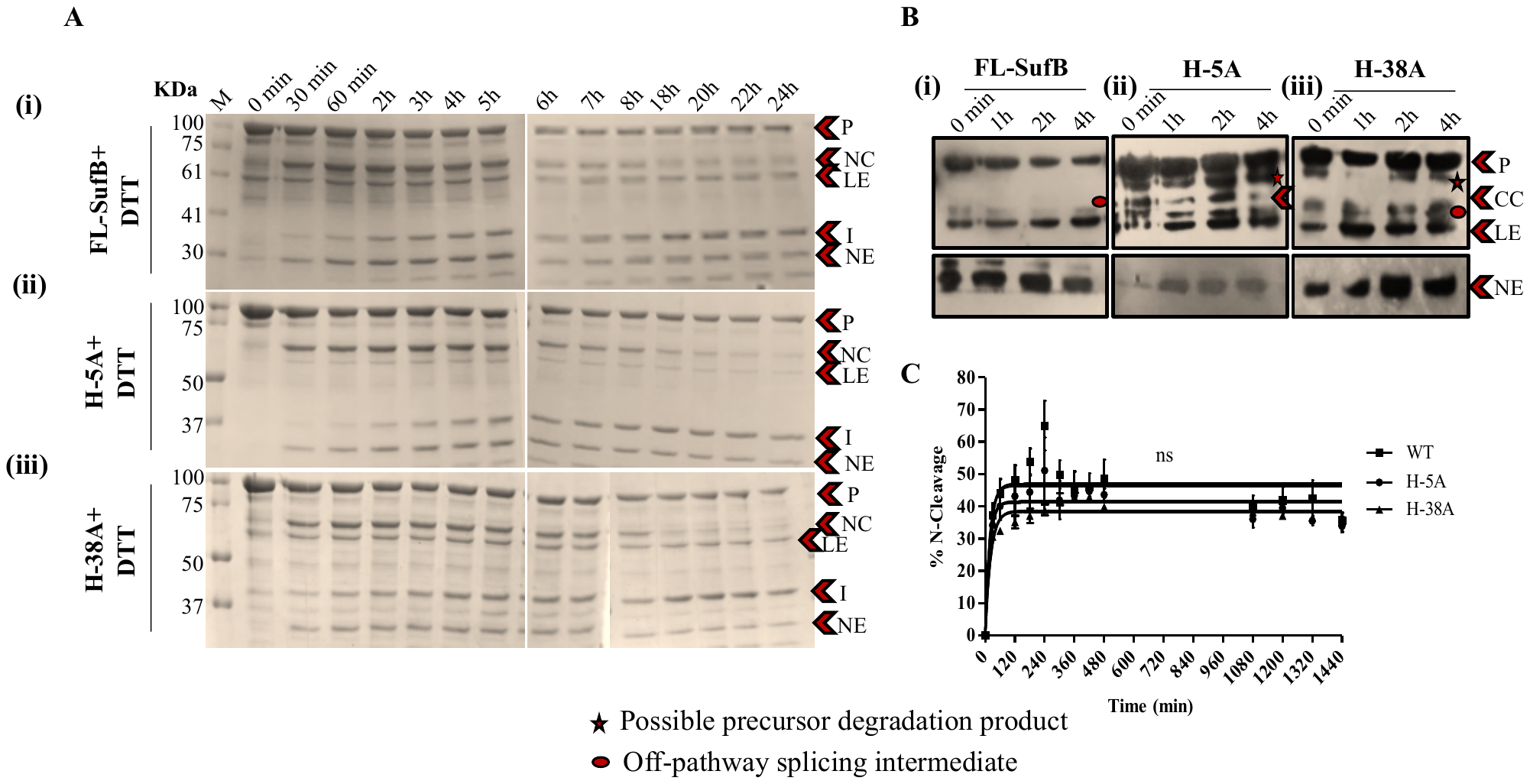

**Figure S3.** (**A**) **Comparison of DTT induced N-terminal cleavage reactions in** (i) FL-SufB precursor, (ii) H-5A and (iii) H-38A mutant SufB proteins. (**B**) Confirmation of identity of splicing and cleavage products for DTT induced N-terminal cleavage assay in (i) FL-SufB precursor, (ii) H-5A and (iii) H-38A SufB mutants via immune blotting. Anti-His antibodies detected presence of 6X(His) tagged P, CC and LE. NE was blotted separately with higher concentration of primary antibody. (**C**) Kinetic Analysis of DTT induced N-terminal cleavage. The cleavage products were calculated and plotted over different time period and the curve was fitted in a pseudo first order reaction, with an equation Y=Y0 + (Plateau-Y0) *(1-exp^(-K*x)^). All the experiments were performed in triplicates and error bars represents (±1) SEM. The comparative analysis shows no statistical significance (p=0.3682). P=Precursor, CC= C-terminal cleavage product, LE=Ligated Extein, NE=N-Extein, M= Protein marker.

**Table S4*. Analysis of different kinetic parameters*** (Ymax, Rate constant and half–life t1/2) for DTT induced N-terminal cleavage reaction in *Mtu* FL-SufB, H-5A and H-38A SufB mutant. These data are extracted from Figure S2C.

| **Kinetic Parameters** | **FL-SufB** | **H-5A** | **H-38A** |
| --- | --- | --- | --- |
| **Ymax** | **46.72±1.7** | **41.54±1.3** | **38.60±0.5** |
| **Rate constant (K)**  **(min^-1^)** | **5.2±0.01x10^-2^** | **5.7±0.017x10^-2^** | **4.4±0.006x10^-2^** |
| **Half-life (t_1/2_ )** | **13.1 min** | **12.05 min** | **15.58 min** |

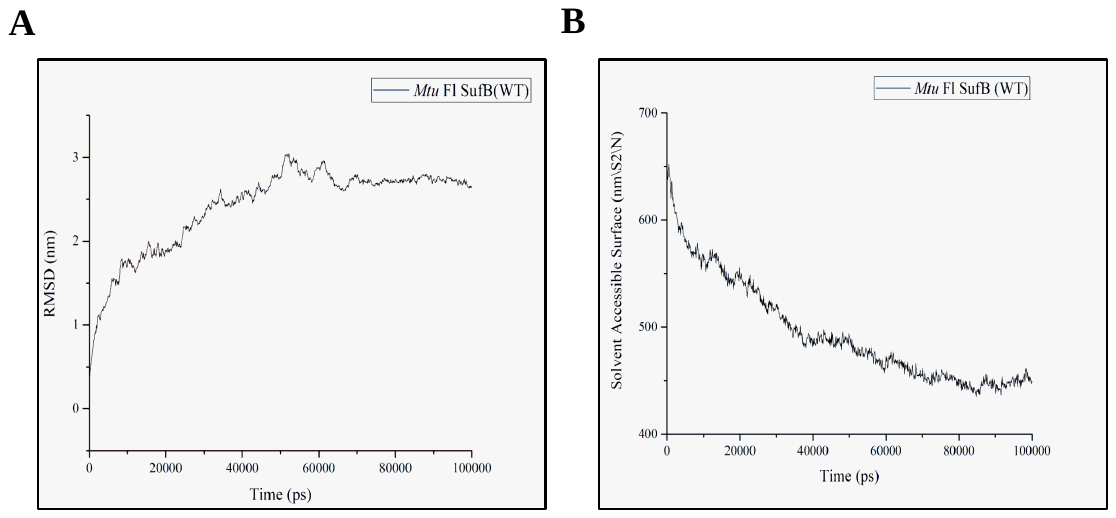

**Figure S4. Analyses on** (**A**) **Root Mean Square Deviation** (RMSD= 2.4±0.5nm) and (**B**) **Solvent Accessible Surface Area** (SASA) of *Mtu* FL-SufB protein after 10^5^ ps (100ns). RMSD reached saturation and there is a minima SASA plot. Therefore, the structure was stabilized after 80ns.

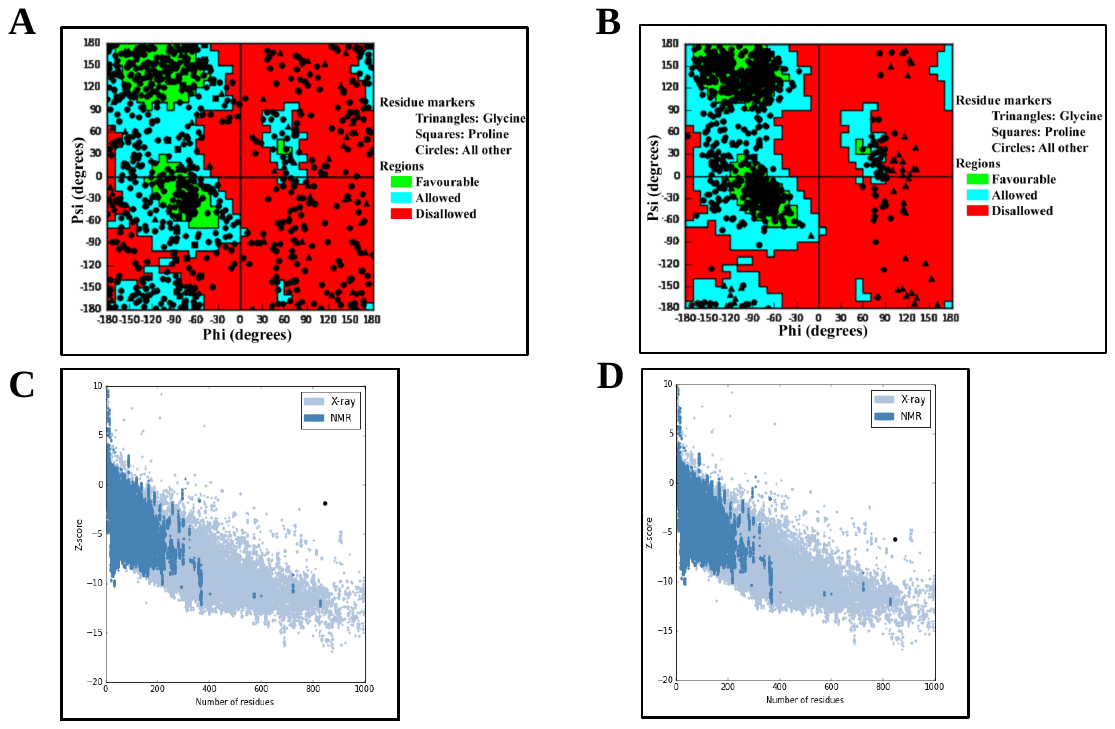

**Figure S5. Model validation of proposed *Mtu* full-length (FL)-SufB precursor structure.** (**A**) Evaluation of homology model by Schrödinger Bio-luminate suite (Ramachandran plot) showing 13% of residues in disallowed region. MD simulation was done for the above mentioned sub-optimized *Mtu* FL-SufB structure in presence of water and NaCl to mimic *in-vivo* condition in GROMACS for proper folding of the protein. (**B**) Ramachandran plot of the optimized *Mtu* FL-SufB structure obtained from the above MD simulation showing only 1% amino acids in the disallowed region. (**C**) ProSA web over all model quality plot of homology model of *Mtu* FL*-*SufB (Z-score -1.86). (**D**) Over all model quality plot of optimized (as mentioned above) *Mtu* Fl-SufB protein (Z-score -5.71) after MD simulation for 100ns.

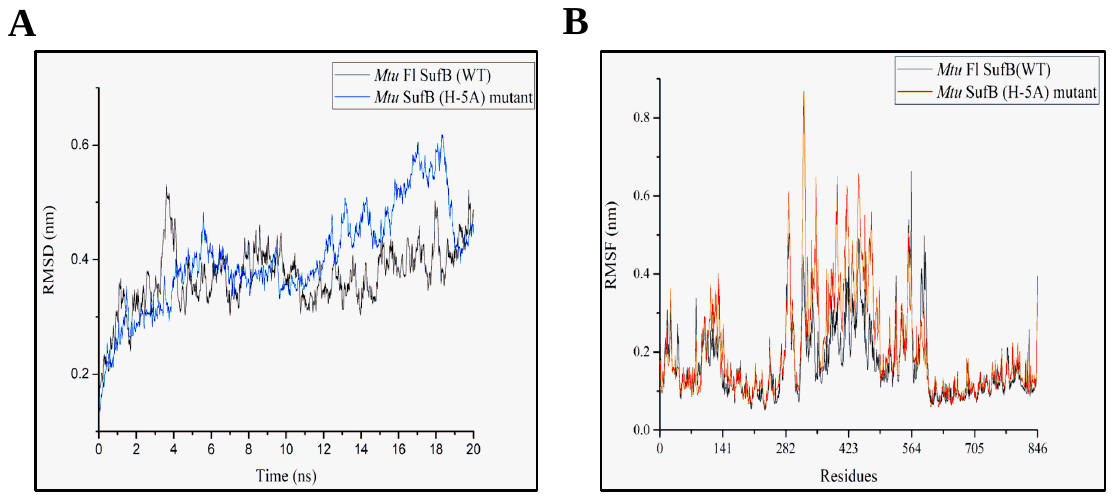

**Figure S6. Analysis on** (**A**) Comparison of Root Mean Square Deviation (RMSD) of FL-SufB and H-5A SufB mutant (RMSD= 3.6±0.5Å and 4.0±0.9Å respectively) simulated for 20ns in aqueous medium. (**B**) Comparison of Root Mean Square Fluctuation (RMSF) of individual amino acid in FL-SufB and H-5A SufB mutant structure.

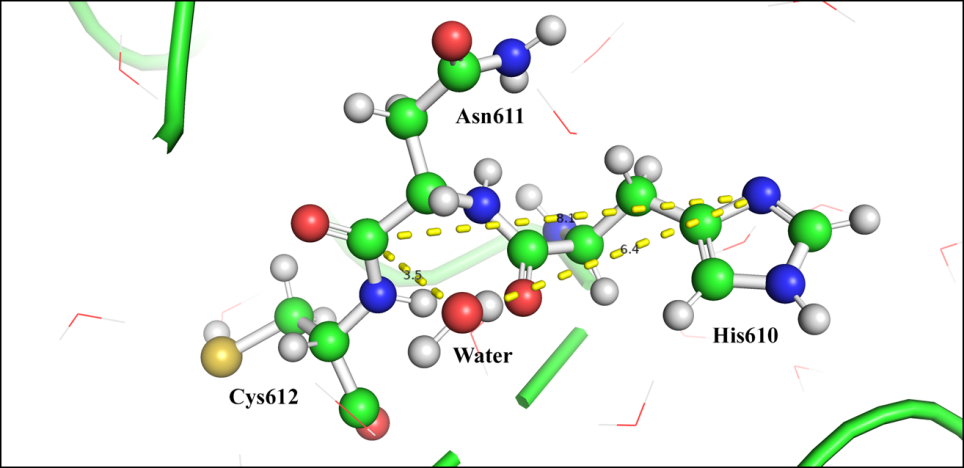

**Figure S7. Structural model showing spatial arrangement of active site residues at C-terminal cleavage junction;** Cys612 (+1), Asn611 (359), His610(358) and a nearby water molecule. Distance (in Å) of possible reacting atoms to peptidyl junction between Cys612 and Asn611 is shown.

**Table S5***.* Molecular Dynamics analysis of *Mtu* FL-Suf B showing average distance between center of mass of active site residues at C-terminal intein~extein junction.

| **Residues** | **Distance in Å(WT)** |
| --- | --- |
| **Cys612~His 610** | **7.39 ± 0.25** |
| **Asn611~His 610** | **4.33±0.25** |
| **Asn611~ Cys612** | **5.52± 0.11** |
